# Tip60-mediated Rheb acetylation links palmitic acid with mTORC1 activation and insulin resistance

**DOI:** 10.1101/2023.08.18.553816

**Authors:** Zengqi Zhao, Qiang Chen, Xiaojun Xiang, Weiwei Dai, Wei Fang, Kun Cui, Baolin Li, Qiangde Liu, Yongtao Liu, Yanan Shen, Yueru Li, Wei Xu, Kangsen Mai, Qinghui Ai

## Abstract

Differences in dietary fatty acid saturation impact glucose homeostasis and insulin sensitivity in vertebrates. Excess dietary intake of saturated fatty acids (SFAs) induces glucose intolerance and metabolic disorders. In contrast, unsaturated fatty acids (UFAs) elicit beneficial effects on insulin sensitivity. However, it remains elusive how SFAs and UFAs signal differentially toward insulin signaling to influence glucose homeostasis. Here, using a croaker model, we report that dietary palmitic acid (PA), but not oleic acid or linoleic acid, leads to dysregulation of mTORC1 signaling which provokes systemic insulin resistance and glucose intolerance. Mechanistically, using croaker primary myocytes, mouse C2C12 myotubes and HEK293T cells, we show that PA-induced mTORC1 activation is dependent on mitochondrial fatty acid β oxidation. Notably, PA profoundly elevates acetyl-CoA derived from mitochondrial fatty acid β oxidation which intensifies Tip60-mediated Rheb acetylation. Subsequently, the induction of Rheb acetylation facilitates hyperactivation of mTORC1 which enhances serine phosphorylation of IRS1 and simultaneously inhibits transcription of IRS1 through impeding TFEB nuclear translocation, leading to impairment of insulin signaling. Furthermore, targeted abrogation of acetyl-CoA produced from fatty acid β oxidation or Tip60-mediated Rheb acetylation by pharmacological inhibition and genetic knockdown rescues PA-induced insulin resistance. Collectively, this study reveals a conserved acetylation-dependent mechanistic insight for understanding the link between fatty acids and insulin resistance, which may provide a potential therapeutic avenue to intervene in the development of T2D.

## Introduction

Insulin resistance, which is considered as a dominant hallmark of type 2 diabetes (T2D)^1^, is related to a variety of metabolic disorders, such as obesity, atherosclerosis and hypertension^2, 3^. The pathogenesis of insulin resistance is associated with genetic mutations including *PPARG*, *IRS1*, *TCF7L2* and *OTUD3*. However, the behavioral and environmental factors can also contribute to insulin resistance in numerous ways^4, 5^. Accumulating evidence have shown that elevated dietary intake of saturated fatty acids (SFAs) is closely correlated with an increased risk of T2D^6, 7^, and palmitic acid (PA), as one of the most abundant circulating SFAs, is also known to trigger the development of insulin resistance^8–12^. On the contrary, dietary intake of unsaturated fatty acids (UFAs) has not been associated with inducing glucose intolerance and may even be beneficial for insulin sensitivity^13^. However, the precise mechanisms underlying the different effects of SFAs and UFAs on insulin signaling and glucose homeostasis are not well established.

Eukaryotes cells have evolved a well-established mechanism to sense the availability of certain nutrients in diet to maintain metabolic homeostasis. mTORC1, which is a central hub of nutrient signaling, integrates a variety of environmental inputs to control cell growth and metabolism^14, 15^. Dysregulation of mTORC1 is associated with a variety of diseases, including T2D, cancer, nonalcoholic fatty liver, Huntington’s disease and Parkinson’s disease^16–18^. The correlation between mTORC1 and T2D is mainly dependent on the existence of several negative feedback mechanisms from mTORC1 and its downstream targets to insulin signaling, which restrain the hyperactivation of mitogenic and anabolism signaling to maintain cellular homeostasis under physiological conditions^19^. Ribosomal S6 kinase (S6K)1, which is a pivotal downstream of mTORC1 has been shown to impair insulin signaling by inducing phosphorylation-dependent degradation of insulin receptor substrate 1 (IRS1) ^20, 21^. Moreover, mTORC1 mediates the phosphorylation and activation of growth factor receptor-bound protein 10 (Grb10), also resulting in the suppression of insulin signaling^22, 23^. Furthermore, imidazole propionate, a metabolite produced by the gut microbiota, is reported to provoke insulin resistance through inducing mTORC1 activation and subsequent phosphorylation and degradation of IRS1^24^. A growing body of evidence has elucidated that mTORC1-induced insulin resistance is related to the phosphorylation and degradation of IRS1. However, whether mTORC1 can affect IRS1 in other manners such as transcriptional regulation under physiological or pathological conditions remains elusive.

As the major regulator of cell growth and metabolism, mTORC1 activity is tightly controlled by a diverse set of upstream signals. Two sets of small G proteins, termed the Rheb and Rag GTPases which integrate the signals from growth factors and nutrients to modulate mTOR kinase activity and intracellular localization respectively, form a center of the regulatory network for mTORC1^25–30^. In addition to the two direct modulations, extensive posttranslational modifications including phosphorylation, acetylation and ubiquitination of mTORC1 components and their associated proteins also influence the activity of mTORC1^31^. Recently, acetylation has been shown to play a vital role in the regulation of mTORC1 activity. Increased acetylation of Raptor by leucine activates mTORC1 signaling which leads to inhibition of autophagy^32, 33^. Consistently, hepatic Acox1 deficiency reduces mTORC1 activity by inhibiting Raptor acetylation under starvation and high fat diet (HFD) conditions in mice^34^. Moreover, the acetylation levels of Rheb are reported to be enhanced during FBS treatment, contributing to the activation of mTORC1^35^. Although the mechanism by which upstream signals such as amino acids regulate mTORC1 has been elucidated, it is still unclear how certain fatty acids modulate mTORC1 activity and whether acetylation modification is involved in the regulation of mTORC1 signaling under fatty acid stimulation.

Teleost fish have evolved conserved systems for nutrient- and pathogen-sensing^36^, which play a central role in maintaining energy homeostasis and resisting pathogenic infection. In addition, our previous work found that teleost fish evolved well-conserved lipid metabolism and acetylation modification systems^37–39^. However, teleost fish have a poor capacity to utilize glucose and are considered to be susceptible to insulin resistance under numerous pathological conditions^40, 41^. Thus, these properties make teleost fish an appropriate model for investigating the pathogenesis of insulin resistance. In this study, using croaker as a *in vivo* model combined with croaker primary myocyte and mouse C2C12 myotube as *in vitro* models, we found that dietary PA-rich diet provokes insulin resistance by inducing hyperactivation of mTORC1. Furthermore, PA-induced mTORC1 activation was dependent on acetyl-CoA derived from mitochondrial fatty acid β oxidation and Tip60-mediated Rheb acetylation. Simultaneously, we also found that mTORC1 could inhibit IRS1 transcription by hindering the nuclear translocation of TFEB, which provided a novel insight into the negative feedback that emanated from mTORC1 to IRS1. Thus, in vertebrates, we demonstrated a conserved acetylation-dependent stress mechanism in response to SFA stimulation, which may provide an attractive strategy to intervene in the development of T2D.

## Results

### PA, but not oleic acid or linoleic acid, induces systemic and cellular insulin resistance

To investigate which fatty acids can provoke insulin resistance, we challenged large yellow croaker with a control (CON), PA rich (PO), oleic acid (OA) rich (OO) or linoleic acid (LA) rich (LO) diet for 10 weeks. There was no significant difference in the final body weight between fish fed CON diet and PO, OO or LO diet (Figures 1A, S1A and S1B). However, the levels of nonesterified free fatty acid (NEFA) in plasma (Figures 1B, S1C and S1D) and triglyceride (TG) in liver (Figures 1C, S1E and S1F) and skeletal muscle (Figures 1D, S1G and S1H) were elevated significantly in fish fed PO, OO or LO diet compared with CON diet, indicating that PA, OA and LA rich feeding may induce aberrant lipid deposition. As abnormal accumulation of intracellular lipids is associated with the development of insulin resistance^42, 43^, we assayed the effect of dietary different fatty acids on glucose homeostasis and insulin sensitivity. Compared with CON diet, PO diet strongly elevated fasting blood glucose levels (Figure 1E) and plasma insulin concentrations (Figure 1F), whereas OA or LA diets had no significant effects (Figures S1I-S1L). Moreover, glucose tolerance (Figure 1G) and insulin tolerance (Figure 1H) were impaired by PO diet, as revealed via the glucose tolerance test (GTT) and the insulin tolerance test (ITT). Furthermore, immunoblotting assays also showed that PO diet, but not OA or LA diet, diminished the phosphorylation levels of AKT in the liver and skeletal muscle (Figures 1I, S1M and S1N), which plays a vital role in insulin signaling. These results suggest that dietary PA leads to systemic insulin resistance in fish, but not OA or LA.

**Figure 1.**
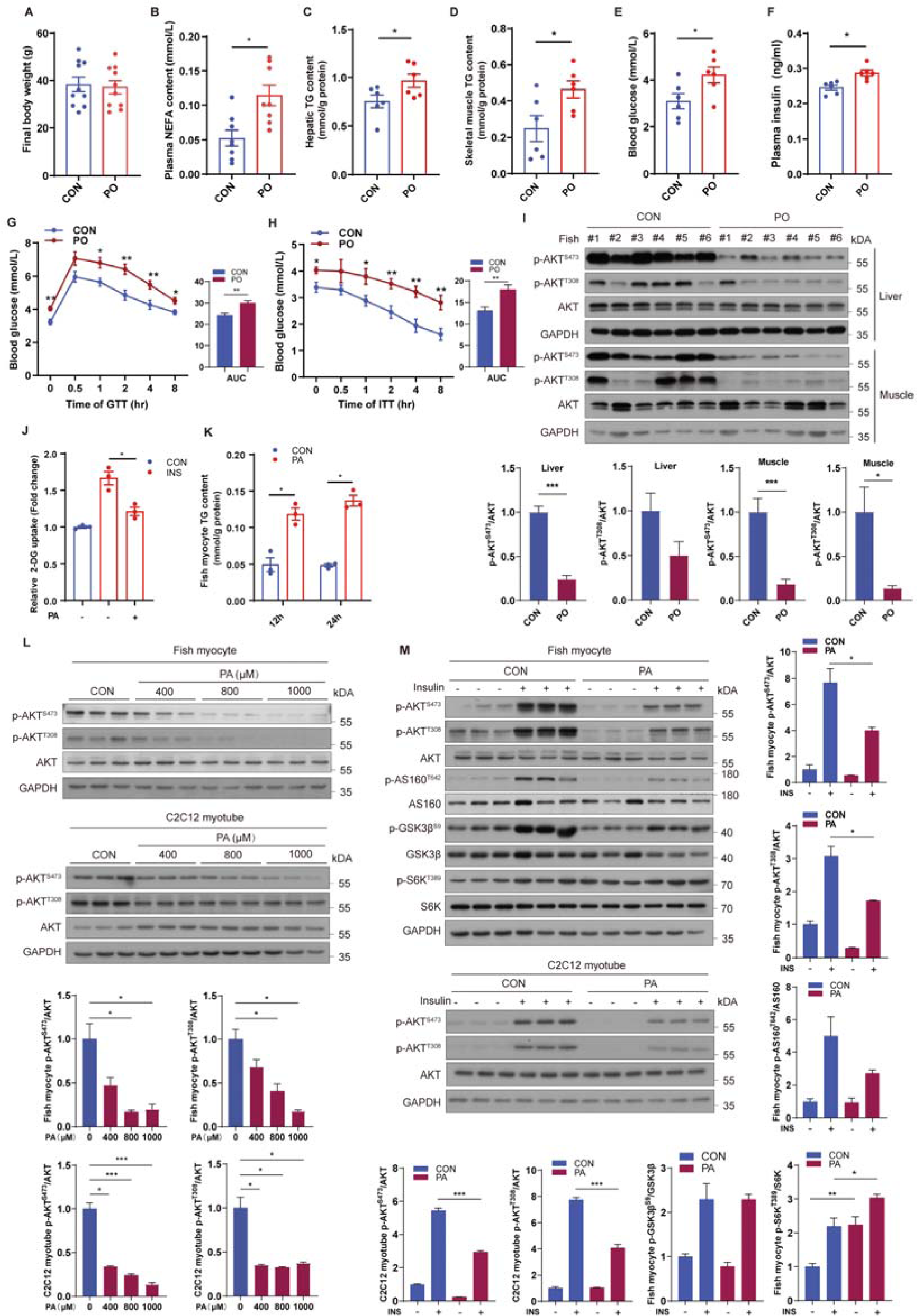
Palmitic acid triggers systemic and cellular insulin resistance. (A-F, I) Fish were fed with control (CON) or palmitic acid (PA) rich (PO) diet for 10 weeks. After 12 h fasting, final body weight and blood glucose were measured; plasma, liver and muscle samples were collected. (A) Final body weight of fish fed CON or PO diet for 10 weeks (n=10). (B) Plasma nonesterified free fatty acid (NEFA) of fish fed different diets (n=8). (C and D) TG levels in liver (C) and skeletal muscle (D) were measured in fish fed different diets (n=6). (E and F) Blood glucose (E) and plasma insulin levels (F) were measured in fasted fish fed different diets (n=6). (G and H) Glucose tolerance (GTT, G) and insulin tolerance tests (ITT, H) were evaluated in fish after treatment with different diets (n=6). (I) Phosphorylation levels of AKT were measured by immunoblotting in liver and skeletal muscle of fish fed different diets (n=6). (J) TG levels in fish myocytes were analyzed under control or PA treatment (n=3). (K) Insulin-stimulated glucose uptake was detected by 2-DG uptake assays under control or PA treatment for 12 h in fish myocytes (n=3). (L) Phosphorylation levels of AKT in fish myocytes and C2C12 myotubes were tested by immunoblotting in the presence of the indicated concentrations of PA for 12 h (n=3). (M) Phosphorylation levels of the indicated proteins in fish myocytes and C2C12 myotubes were detected by immunoblotting (n=3). Cells were pretreated with control or PA for 12 h, and then stimulated with insulin for 5 min. The results are presented as the mean ± SEM and were analyzed using independent *t*-tests (\**p* < 0.05, \*\**p* < 0.01, \*\*\**p* < 0.001). See also Figure S1.

To further investigate the role of PA in glucose homeostasis and insulin sensitivity, fish myocytes and mouse differentiated C2C12 myotubes were treated with PA. In line with the *in vivo* results, PA treatment increased the TG levels of fish myocytes (Figure 1J). Moreover, glucose uptake stimulated by insulin was inhibited under PA treatment in fish myocytes (Figure 1K), as unveiled by 2-deoxy-D-glucose (2-DG) uptake assays. Likewise, PA treatment reduced the phosphorylation levels of AKT in a dose-dependent manner (Figure 1L) and impeded insulin from boosting the phosphorylation of AKT in fish myocytes and C2C12 myotubes (Figure 1M). To determine whether OA or LA can impair cellular insulin signaling, we also treated cells with OA or LA but found that both treatments failed to inhibit glucose uptake (Figures S1O and S1P) and phosphorylation of AKT with or without insulin stimulation (Figures S1Q-S1T). Together, these results indicate that PA provokes systemic and cellular insulin resistance, whereas OA or LA had no effect on glucose homeostasis and insulin sensitivity.

### Hyperactivation of mTORC1 contributes to PA-induced insulin resistance

To elucidate the underlying mechanism of insulin resistance induced by PA, we assayed the activities of AKT downstream signaling pathways under PA treatment. Notably, unlike other AKT downstream substrates (GSK3β and AS160), PA treatment failed to decrease the levels of phosphorylated S6K, an indicator of mTORC1 activity, under insulin stimulation (Figure 1M). Considering that mTORC1 can lead to feedback inhibition of insulin signaling^44, 45^, we speculated that mTORC1 is involved in PA-induced insulin resistance. To confirm this hypothesis, we therefore estimated the effect of PO diet on mTORC1 signaling and found that the phosphorylation levels of S6K and S6 were strongly promoted in skeletal muscle of fish fed PO diet (Figure 2A). Moreover, in cultured cells, PA treatment elevated the phosphorylation levels of S6K and S6 in a time- and dose-dependent manner (Figures 2B and 2C), revealing that PA can provoke hyperactivation of mTORC1.

**Figure 2.**
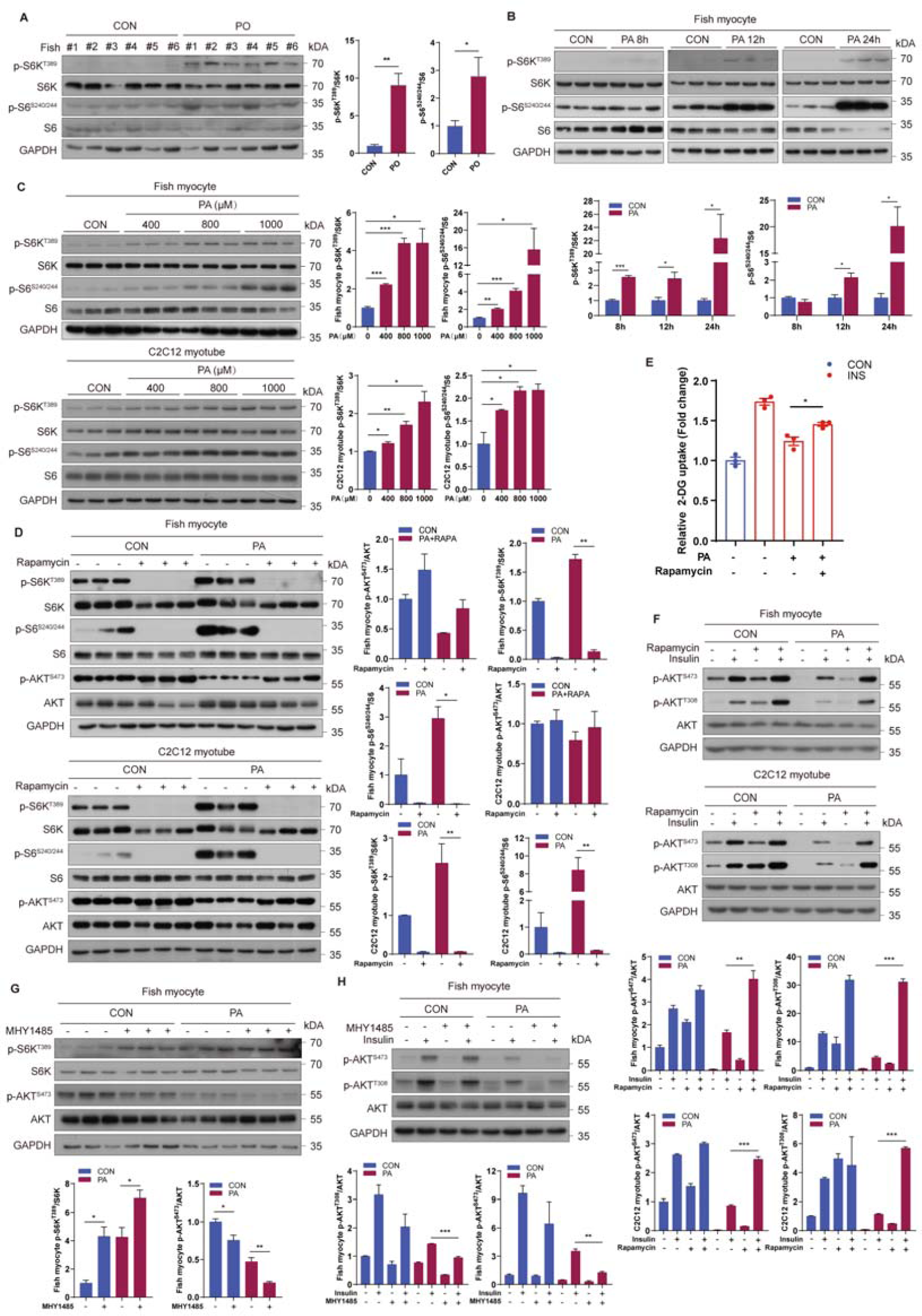
Hyperactivation of mTORC1 is associated with PA-induced insulin resistance. (A) mTORC1 pathway activity was measured by immunoblotting for the phosphorylation of S6K and S6 in skeletal muscle of fish fed CON or PO diet (n=6). (B) mTORC1 pathway activity was tested by immunoblotting in fish myocytes under control or PA treatment for 8 h, 12 h and 24 h (n=3). (C) mTORC1 pathway activity was assayed by immunoblotting in fish myocytes and C2C12 myotubes in the presence of the indicated concentrations of PA for 12 h (n=3). (D) mTORC1 pathway activity was analyzed by immunoblotting in fish myocytes and C2C12 myotubes under control or PA treatment in the presence or absence of rapamycin for 12 h (n=3). (E) Insulin-stimulated glucose uptake was detected by 2-DG uptake assays in fish myocytes under control or PA treatment with or without rapamycin for 12 h (n=3). (F) Phosphorylation levels of AKT in fish myocytes and C2C12 myotubes were detected by immunoblotting (n=3). Cells were pretreated with control or PA treatment in the presence or absence of rapamycin for 12 h, and then stimulated with insulin for 5 min. (G) mTORC1 pathway activity was measured by immunoblotting in fish myocytes under control or PA treatment with or without MHY1485 for 12 h (n=3). (H) Phosphorylation levels of AKT in fish myocytes were tested by immunoblotting (n=3). Cells were pretreated with control or PA treatment in the presence or absence of MHY1485 for 12 h, and then stimulated with insulin for 5 min. The results are presented as the mean ± SEM and were analyzed using independent *t*-tests (\**p* < 0.05, \*\**p* < 0.01, \*\*\**p* < 0.001). See also Figure S2.

To further investigate the role of mTORC1 during PA-induced insulin resistance, fish myocytes and C2C12 myotubes were incubated with rapamycin, a potent mTORC1 inhibitor. The results showed that rapamycin treatment indeed prevented PA-induced mTORC1 activation (Figure 2D). Meanwhile, the suppression of insulin stimulated-glucose uptake by PA treatment was restored upon rapamycin treatment (Figure 2E). Furthermore, insulin-intensified phosphorylation of AKT was improved when mTORC1 was inhibited by rapamycin (Figure 2F). These results indicated that PA-induced insulin resistance is associated with the activation of mTORC1. Further confirming this notion, fish myocytes were treated with the mTORC1 activator MHY1485. As expected, MHY1485 treatment enhanced mTORC1 activity (Figure 2G) and restrained insulin from boosting the phosphorylation of AKT in the presence or absence of PA (Figure 2H). Given that OA or LA did not alter glucose homeostasis and insulin sensitivity, we also assayed the effect of OA or LA on mTORC1 signaling. The results showed that OA and LA failed to induce mTORC1 activity *in vivo* (Figure S2A and S2B) and *in vitro* (Figure S2C and S2D). These results suggest that the difference in mTORC1 regulation among PA, OA or LA may lead to divergent effects on insulin signaling, and that also further illustrate the relevance between activation of mTORC1 and diet-induced insulin resistance.

### Mitochondrial fatty acid β oxidation is required for PA-induced mTORC1 activation and insulin resistance

The above results showed that the induction of mTORC1 signaling occurred after 8 h of PA stimulation (Figure 2B). Thus, we speculated that activation of mTORC1 by PA may not be related to its role as a signaling molecule but dependent on its metabolic pathway. Considering that fatty acids must first form fatty acid-CoA in order for anabolism or catabolism to proceed, we detected the contents of acyl-CoA and acylcarnitine using LC–MS. The results showed that PA treatment increased the contents of short/medium-chain acyl-CoA and acylcarnitine (Figures 3A and S3A), indicating that PA may induce mitochondrial fatty acid β oxidation. Moreover, seahorse real-time cell metabolic analysis showed that PA treatment enhanced mitochondrial OCR and elevated maximal oxygen consumption rates compared with OA or LA treatment (Figure 3B). Similarly, in comparison with CON diet, fish fed PO diet exhibited higher mRNA expression levels of fatty acid β oxidation-related genes in muscle (Figure 3C). However, OO or LO diet failed to increase mRNA levels of fatty acid β oxidation-related genes (Figure S3B and S3C). Furthermore, compared with OA or LA treatment, PA-induced increase of fatty acid oxidation gene expressions was more robust *in vitro* (Figures 3D, 3E, S3D and S3E). Thus, these results revealed that compared with OA or LA, PA may be preferred to enter the mitochondria for fatty acid β oxidation and that PA-induced mTORC1 activation is associated with mitochondrial fatty acid β oxidation possibly.

**Figure 3.**
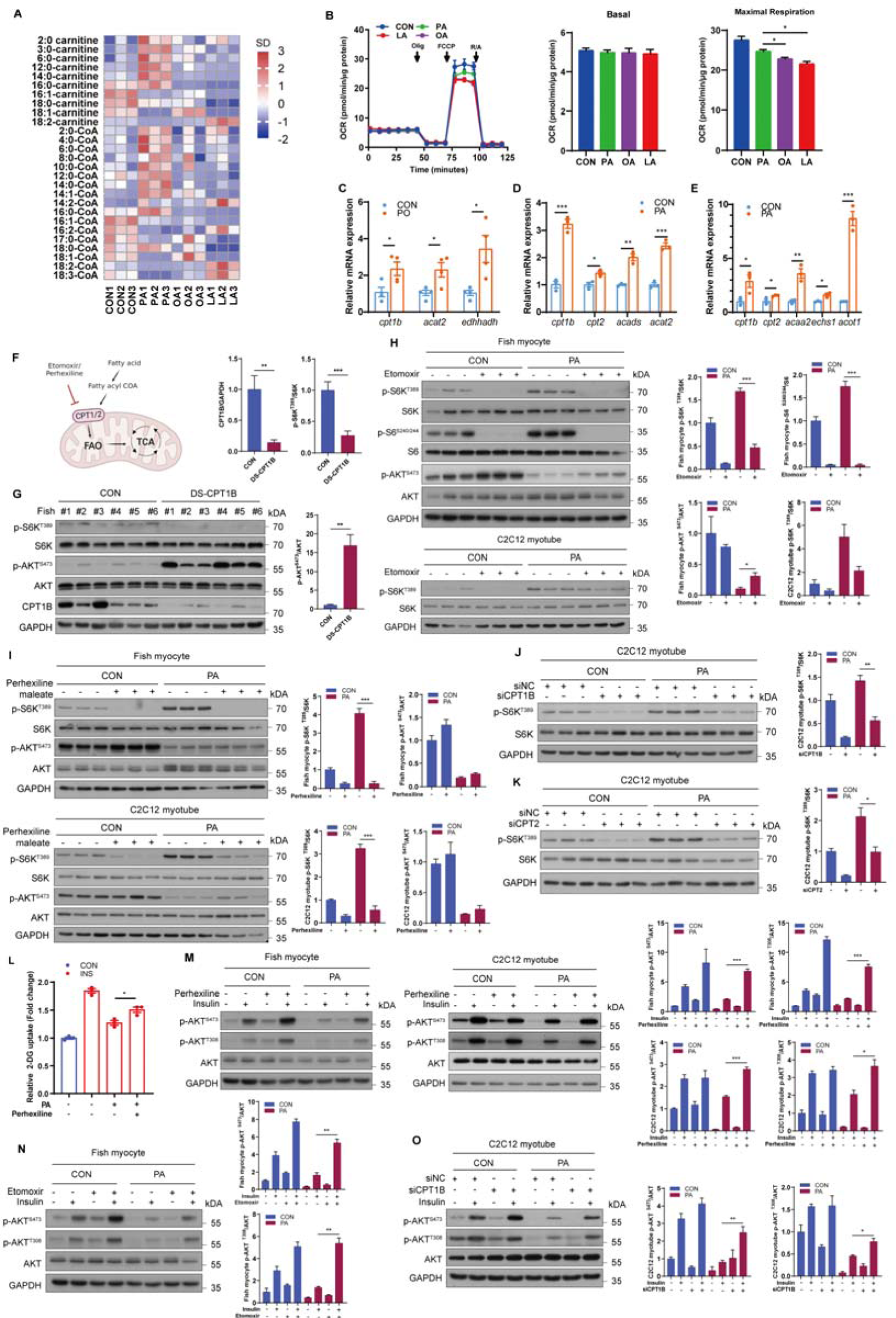
Mitochondrial fatty acid β oxidation is required for PA-induced mTORC1 activation and insulin resistance. (A) Heat map of the contents of acyl-CoAs and acyl-carnitines in fish myocytes treated with PA, OA or LA for 12h. The relative fold change for each factor in each sample is represented as the relative average increase (red) or decrease (blue) (n=3). (B) Oxygen consumption rate (OCR) traces as measured by seahorse XF 24 Flux Analyzer in fish myocytes treated with PA, OA or LA for 12h (n=3). (C) Relative mRNA levels of mitochondrial fatty acid β oxidation-related genes were analyzed by quantitative PCR in the muscle of fish fed CON or PO diet (n=4). (D and E) Relative mRNA levels of mitochondrial fatty acid β oxidation-related genes were measured by quantitative PCR in fish myocytes (B) and C2C12 myotubes (C) with control or PA treatment for 12 h (n=3). (F) Schematic representation of the main cellular pathways involved in mitochondrial fatty acid β oxidation. (G) The activities of mTORC1 and AKT were analyzed via immunoblotting in fish with intraperitoneal injection of control dsRNA or dsRNA targeting CPT1B for 36 h (n=6). (H) The activities of mTORC1 and AKT were assayed by immunoblotting in fish myocytes and C2C12 myotubes treated with control or etomoxir treatment in the presence or absence of PA for 12 h (n=3). (I) The activities of mTORC1 and AKT were assayed by immunoblotting in fish myocytes and C2C12 myotubes treated with control or perhexiline maleate treatment in the presence or absence of PA for 12 h (n=3). (J) The activity of mTORC1 signaling was examined by immunoblotting in C2C12 myotubes transfected with control siRNA or siRNA against CPT1B in the presence or absence of PA (n=3). (K) The activity of mTORC1 signaling was measured by immunoblotting in C2C12 myotubes transfected with control siRNA or siRNA against CPT2 in the presence or absence of PA (n=3). (L) Insulin-stimulated glucose uptake was detected by 2-DG uptake assays in fish myocytes under control or perhexiline maleate in the presence or absence of PA for 12 h (n=3). (M) AKT phosphorylation levels in fish myocytes and C2C12 myotubes were tested by immunoblotting (n=3). Cells were pretreated with control or perhexiline maleate treatment in the presence or absence of PA for 12 h, and then stimulated with insulin for 5 min. (N) AKT phosphorylation levels in fish myocytes were tested by immunoblotting (n=3). Cells were pretreated with control or etomoxir treatment in the presence or absence of PA for 12 h, and then stimulated with insulin for 5 min. (O) AKT phosphorylation levels in C2C12 myotubes were measured by immunoblotting (n=3). Cells were transfected with control siRNA or siRNA against CPT1B and pretreated with control or PA treatments for 12 h, and then stimulated with insulin for 5 min. The results are presented as the mean ± SEM and were analyzed using independent *t*-tests (\**p* < 0.05, \*\**p* < 0.01, \*\*\**p* < 0.001). See also Figure S3.

Considering that CPT1B and CPT2 are rate-limiting enzymes of fatty acid β oxidation in muscle, we suppressed CPT1B and CPT2 by pharmacological inhibition and genetic knockdown to block fatty acid oxidation (Figure 3F). The results showed that intraperitoneal injection of dsCPT1B in fish strongly reduced the expression of CPT1B and inhibited the activity of mTORC1 in muscle (Figure 3G). Moreover, respectively incubating fish myocytes and C2C12 myotubes with etomoxir or perhexiline maleate, two potent CPT1 inhibitors, attenuated the induction of mTORC1 signaling under PA treatment (Figures 3H and 3I). Similarly, CPT1B and CPT2 knockdown by small interfering RNA (siRNA) (Figures S3F-S3I) also abrogated the activation of mTORC1 under PA treatment in C2C12 myotubes (Figures 3J and 3K). These results suggest that PA-provoked mTORC1 activation is dependent on mitochondrial fatty acid β oxidation and that the different effects of fatty acid β oxidation among PA, OA or LA may contribute to divergent effects on mTORC1 regulation.

Given that hyperactivation of mTORC1 signaling may account for insulin resistance, we validated whether the inhibition of mitochondrial fatty acid β oxidation contributed to the recovery of insulin sensitivity under PA treatment. As expected, the suppression of insulin-stimulated glucose uptake by PA treatment was relieved in fish myocytes with pharmacological inhibition of fatty acid β oxidation (Figure 3L). Consistently, the phosphorylation levels of AKT were enhanced in the muscle of fish with intraperitoneal injection of dsCPT1B (Figure 3G). Furthermore, inhibition of CPT1 by pharmacological inhibitors or siRNA knockdown also improved the suppression of insulin increased phosphorylation of AKT under PA treatment in fish myocytes and C2C12 myotubes (Figures 3M-3O). Collectively, these results demonstrate that mitochondrial fatty acid β oxidation plays a vital role in PA-induced mTORC1 activation and subsequent insulin resistance.

### Acetyl-CoA derived from mitochondrial fatty acid β oxidation enhances Rheb acetylation to boost mTORC1 activation and insulin resistance

As acetyl-CoA is the major and final metabolite of mitochondrial fatty acid β oxidation^46^, we investigated whether fatty acid β oxidation produced acetyl-CoA mediated the induction of mTORC1 signaling and insulin resistance. Compared with CON diet, PO diet strongly increased acetyl-CoA levels in muscle (Figure 4A). Likewise, PA treatment significantly elevated the levels of intracellular acetyl-CoA in a dose-dependent manner in fish myocytes (Figure 4B). Moreover, fatty acid β oxidation blocked by perhexiline maleate diminished the induction of acetyl-CoA levels under PA treatment (Figure 4C). Furthermore, we directly measured the contribution of PA to the total cellular acetyl-CoA pool using metabolic flux assays. The results showed that exceeding 60% of the acetyl-CoA pool was PA derived and exceeding 50% of the other short/medium-chain fatty acid-CoA pools were PA derived (Figure 4D). Thus, these results indicated that acetyl-CoA derived from mitochondrial fatty acid β oxidation may play a potential role in PA-induced hyperactivation of mTORC1. Given that ATP citrate lyase (ACLY) governs “citrate transport” and is responsible for acetyl-CoA transfer from mitochondria (Figure 4E), dsRNA-mediated ACLY knockdown was performed *in vivo* and the results showed that dsACLY decreased the activity of mTORC1 in muscle (Figure 4F). Similarly, inhibition of ACLY with BMS-303141 restrained the induction of mTORC1 under PA treatment in fish myocytes (Figure 4G) and ACLY knockdown by siRNA (Figures S4A and S4B) also abrogated PA-induced mTORC1 activation in C2C12 myotubes (Figures 4H). Furthermore, sodium acetate treatment which can enhance acetyl-CoA levels via acetyl-CoA synthetase 2 (ACSS2), promoted mTORC1 activation in a dose-dependent manner in fish myocytes and C2C12 myotubes (Figure 4I). Notably, impaired PA-induced mTORC1 activation by the inhibition of fatty acid β oxidation was rescued by sodium acetate treatment in fish myocytes and C2C12 myotubes (Figure 4J). These results indicated that PA-provoked mTORC1 activation is dependent on the acetyl-CoA derived from mitochondrial fatty acid β oxidation.

**Figure 4.**
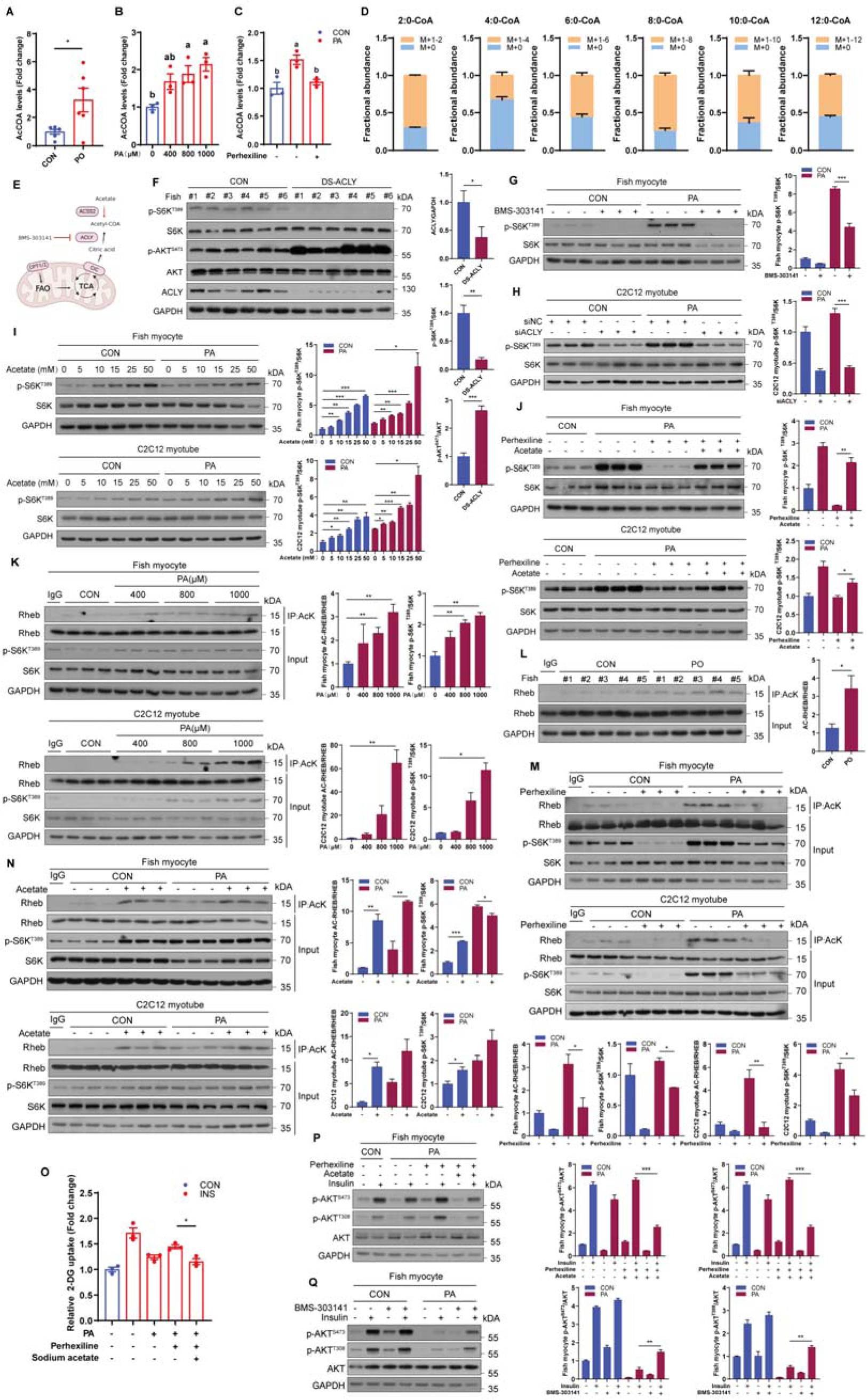
Acetyl-CoA derived from mitochondrial fatty acid β oxidation triggers mTORC1 activation and insulin resistance through enhancing Rheb acetylation. (A) Acetyl-CoA levels were measured in the muscle of fish fed CON or PO diet (n = 6). (B) Acetyl-CoA levels in fish myocytes were measured in the presence of the indicated concentrations of PA for 12 h (n = 3). (C) Acetyl-CoA levels were measured in fish myocytes under control or PA treatment with or without perhexiline maleate for 12 h (n = 3). (D) Acetyl-CoAs labeling pattern from fish myocytes treated with [U-^13^C_16_]-labeled palmitate (n = 5). (E) Schematic representation of the main cellular pathways involved in ACLY-governed citrate transport and ACSS2-mediated acetyl-CoA production. (F) The activities of mTORC1 and AKT were assayed via immunoblotting in fish with intraperitoneal injection of control dsRNA or dsRNA targeting ACLY for 36 h (n=6). (G) The activity of mTORC1 was tested by immunoblotting in fish myocytes with control or BMS-303141 treatment in the presence or absence of PA for 12 h (n=3). (H) The activity of mTORC1 signaling was analyzed by immunoblotting in C2C12 myotubes transfected with control siRNA or siRNA against ACLY under control or PA treatment (n=3). (I) The activity of mTORC1 signaling was measured by immunoblotting in fish myocytes and C2C12 myotubes with the indicated concentrations of sodium acetate under control or PA treatment for 12 h (n=3). (J) The activity of mTORC1 signaling was measured by immunoblotting in fish myocytes and C2C12 myotubes under control or perhexiline maleate treatment with or without sodium acetate addition in the presence or absence of PA for 12 h (n=3). (K) Immunoblotting of Rheb acetylation and S6K phosphorylation in fish myocytes and C2C12 myotubes treated with the indicated concentrations of PA for 12 h (n=3). (L) Immunoblotting of Rheb acetylation in the muscle of fish fed CON or PO diet (n = 5). (M) Immunoblotting of Rheb acetylation and S6K phosphorylation in fish myocytes and C2C12 myotubes treated with control or perhexiline maleate treatment in the presence or absence of PA for 12 h (n=3). (N) Immunoblotting of Rheb acetylation and S6K phosphorylation in fish myocytes and C2C12 myotubes treated with the indicated concentrations of sodium acetate in the presence or absence of PA for 12 h (n=3). (O) Insulin-stimulated glucose uptake was detected by 2-DG uptake assays in fish myocytes under control or perhexiline maleate treatment with or without sodium acetate addition in the presence or absence of PA for 12 h (n=3). (P) AKT phosphorylation levels were tested by immunoblotting in fish myocytes (n=3). Cells were pretreated under control or perhexiline maleate treatment with or without sodium acetate addition in the presence or absence of PA for 12 h, and then stimulated with insulin for 5 min. (Q) AKT phosphorylation levels were detected by immunoblotting in fish myocytes (n=3). Cells were pretreated with control or BMS-303141 treatment in the presence or absence of PA for 12 h, and then stimulated with insulin for 5 min. The results are presented as the mean ± SEM and were analyzed using independent *t*-tests (\**p* < 0.05, \*\**p* < 0.01, \*\*\**p* < 0.001) and Tukey’s tests (bars bearing different letters are significantly different among treatments (*p* < 0.05)). See also Figure S4.

Considering that acetyl-CoA is the direct acetyl donor of acetylation^47^ and that upregulation of Rheb or Raptor acetylation contributes to mTORC1 activation^48^, we measured the acetylation levels of Rheb and Raptor under PA treatment to further investigate the exact mechanism of acetyl-CoA mediated-mTORC1 activation. PA treatment elevated the acetylation levels of Rheb in a dose-dependent manner (Figure 4K), but had no effect on the acetylation levels of Raptor (Figures S4C and S4D) in fish myocytes and C2C12 myotubes. Furthermore, acetylation levels of Rheb were elevated in the muscle of fish fed PO diet compared with CON diet (Figure 4L). Consistently, inhibition of fatty acid β oxidation by perhexiline maleate attenuated PA-stimulated Rheb acetylation (Figure 4M). However, fish myocytes and C2C12 myotubes with sodium acetate addition exhibited enhanced Rheb acetylation (Figure 4N). Together, these results suggest that acetyl-CoA produced by fatty acid β oxidation activates mTORC1 signaling through increasing Rheb acetylation.

Next, we investigated the role of acetyl-CoA in PA-induced insulin resistance. As expected, the phosphorylation levels of AKT were enhanced in the muscle of fish with dsACLY injection (Figure 4F) and sodium acetate addition blocked the recovery of insulin-stimulated glucose uptake and phosphorylation levels of AKT by perhexiline maleate under PA condition (Figures 4O and 4P). Moreover, inhibition of ACLY with BMS-303141 promoted insulin-stimulated phosphorylation levels of AKT in the presence of PA (Figure 4Q). These results indicate that acetyl-CoA derived from fatty acid β oxidation mainly mediates PA-induced mTORC1 activation and insulin resistance.

### Tip60 regulates mTORC1 activity and insulin sensitivity through acetylating Rheb under PA treatment

As an increase in acetyl-CoA can activate lysine acetyltransferases (KATs) which are responsible for protein acetylation^49^, we further identified which KATs mediated the acetylation of Rheb under PA treatment. The mRNA expression levels of *cbp*, *gcn5* and *tip60* were elevated in fish myocytes and C2C12 myotubes under PA treatment (Figures 5A and 5B). Therefore, inhibitors of CBP/P300 (C646 and spermidine), GCN5 (MB-3) and Tip60 (MG149) were used to evaluate whether these KATs mediated the regulation of mTORC1 signaling under PA condition. Inhibiting CBP/P300 or GCN5 did not reduce the activity of mTORC1 under PA condition in C2C12 myotubes (Figures S5A-S5C). In contrast, treated fish myocytes and C2C12 myotubes with the Tip60 inhibitor MG149 prevented the induction of mTORC1 activity under PA treatment (Figure 5C), and Tip60 knockdown by siRNA also blocked PA-induced mTORC1 activation in C2C12 myotubes (Figures 5D, S5D and S5E), suggesting that Tip60 may be involved in the regulation of mTORC1 activity under PA treatment. To further investigate whether the regulation of mTORC1 by Tip60 is dependent on the acetylation of Rheb, the interaction between Tip60 and Rheb was analyzed via co-immunoprecipitation assays. The results showed that Tip60 can interact with Rheb in HEK293T cells (Figure 5E). Moreover, overexpressed Tip60 reinforced the acetylation of Rheb and phosphorylation levels of S6K in HEK293T cells (Figure 5F). Furthermore, pharmacological inhibition of Tip60 by MG149 treatment or Tip60 knockdown by siRNA also impaired PA-induced acetylation of Rheb (Figures 5G and 5H). Notably, OA and LA treatments had no effect on the mRNA expression of *tip60*, further confirming the role of Tip60 in the regulation of mTORC1 signaling (Figures S5F and S5G). These results strongly supported the notion that Tip60 mediated the acetylation of Rheb under PA treatment.

**Figure 5.**
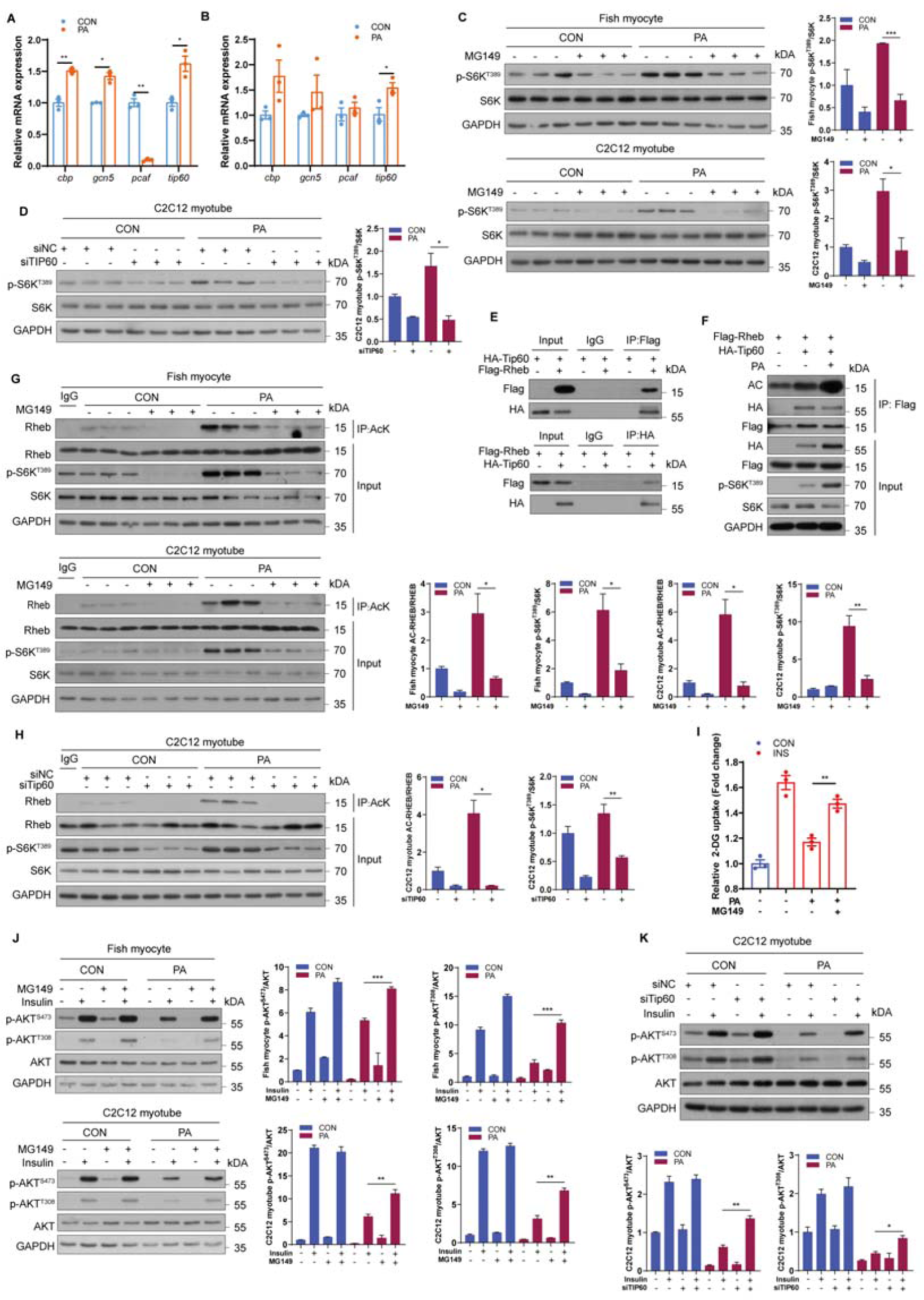
Tip60 regulates mTORC1 activity and insulin sensitivity through acetylating Rheb under PA treatment. (A) Relative mRNA levels of acetyltransferase genes (*cbp*, *gcn5*, *pcaf* and *tip60*) were analyzed by quantitative PCR in fish myocytes under control or PA treatment (n=3). (B) Relative mRNA levels of acetyltransferase genes (*cbp*, *gcn5*, *pcaf* and *tip60*) were analyzed by quantitative PCR in C2C12 myotubes with control or PA treatment (n=3). (C) Immunoblotting of S6K phosphorylation in fish myocytes and C2C12 myotubes treated with control or MG149 treatment in the presence or absence of PA for 12 h (n=3). (D) The activity of mTORC1 signaling was measured by immunoblotting in C2C12 myotubes transfected with control siRNA or siRNA against Tip60 under control or PA treatment (n=3). (E) HEK293T cells were transfected with plasmids as indicated, and the protein was extracted for co-immunoprecipitation (Co-IP) to assay the interaction between Rheb and Tip60. (F) HEK293T cells were transfected with FLAG-Rheb together with or without HA-Tip60 plasmids in the presence or absence of PA, and acetylation levels of Rheb and phosphorylation levels of S6K were measured via immunoblotting. (G) Immunoblotting of Rheb acetylation and S6K phosphorylation in fish myocytes and C2C12 myotubes treated with control or MG149 treatment in the presence or absence of PA for 12 h (n=3). (H) Immunoblotting of Rheb acetylation and S6K phosphorylation in C2C12 myotubes transfected with control siRNA or siRNA against Tip60 under control or PA treatment (n=3). (I) Insulin-stimulated glucose uptake was measured by 2-DG uptake assays in fish myocytes treated with control or MG149 in the presence or absence of PA for 12 h (n=3). (J) AKT phosphorylation levels were assayed by immunoblotting in fish myocytes and C2C12 myotubes (n=3). Cells were pretreated with control or MG149 treatment in the absence or presence of PA for 12 h, and then stimulated with insulin for 5 min. (K) AKT phosphorylation levels were assayed by immunoblotting in C2C12 myotubes (n=3). Cells were transfected with control siRNA or siRNA against Tip60 and pretreated with control or PA for 12 h, and then stimulated with insulin for 5 min. The results are presented as the mean ± SEM and were analyzed using independent *t*-tests (\**p* < 0.05, \*\**p* < 0.01, \*\*\**p* < 0.001). See also Figure S5.

Since the above results demonstrated that Rheb acetylation is associated with insulin resistance in PA condition, we explored whether Tip60 was a potential therapeutic target for insulin resistance. Inhibition of Tip60 by MG149 attenuated PA-induced suppression of insulin-stimulated glucose uptake in C2C12 myotubes (Figure 5I). Consistently, inhibition of Tip60 by MG149 treatment or siRNA knockdown restored insulin-stimulated phosphorylation levels of AKT in the presence of PA (Figures 5J and 5K). Together, these results suggest that Tip60, which mediates the regulation of Rheb acetylation, may be a novel therapeutic target for insulin resistance.

### Negative regulation of IRS1 by mTORC1 is associated with PA-induced insulin resistance

To further investigate the mechanism of insulin resistance induced by mTORC1 activation under PA treatment, we measured the effect of PA on IRS1 phosphorylation, considering that mTORC1 signaling is reported to provoke IRS1 serine phosphorylation^50, 51^. The results showed that PO diet intensified the S636/S639 phosphorylation in fish muscle, compared with CON diet (Figure 6A). In line with the *in vivo* results, PA treatment elevated S636/S639 phosphorylation but decreased the Y612 phosphorylation of IRS1 in a dose-dependent manner in fish myocytes and C2C12 myotubes (Figure 6B). Moreover, incubating fish myocytes and C2C12 myotubes with rapamycin or Torin1 (mTOR inhibitor) blocked the increase of IRS1 S636/S639 phosphorylation levels induced by PA treatment (Figure 6C), and MHY1485 treatment aggravated the induction of IRS1 S636/S639 phosphorylation levels under PA treatment (Figure 6D). Thus, these results indicated that PA-induced alteration of IRS1 phosphorylation which may contribute to insulin resistance is dependent on mTORC1 signaling. Intriguingly, we observed that the protein levels of IRS1 were reduced under PA treatment (Figures 6A and 6B), raising the possibility that PA may impede the transcription of IRS1. Therefore, we further detected the mRNA levels of *irs1* under PA condition and found that fish fed PO diet had significantly lower mRNA levels of *irs1* in muscle compared with CON diet (Figure 6E). Likewise, PA treatment strongly reduced the mRNA levels of *irs1* in fish myocytes and C2C12 myotubes (Figures 6F and 6G). Nevertheless, PA had no effect on the mRNA levels of *insr* and *irs2 in vivo* and *in vitro* (Figures 6E-6G). Furthermore, rapamycin and Torin1 treatments attenuated the decrease of *irs1* mRNA levels under PA treatment in fish myocytes and C2C12 myotubes (Figure 6H). These results indicate that PA-induced mTORC1 activation contributes to the inhibition of IRS1 transcription. Given the role of IRS1 in insulin signaling, we also analyzed the expression of IRS1 under OA and LA treatments. The results showed that OA or LA treatment failed to reduce the mRNA levels of *irs1 in vivo* (Figure S6A) and *in vitro* (Figure S6B). Collectively, these results imply that alteration of IRS1 phosphorylation and transcription by mTORC1 is involved in the insulin resistance induced by PA.

**Figure 6.**
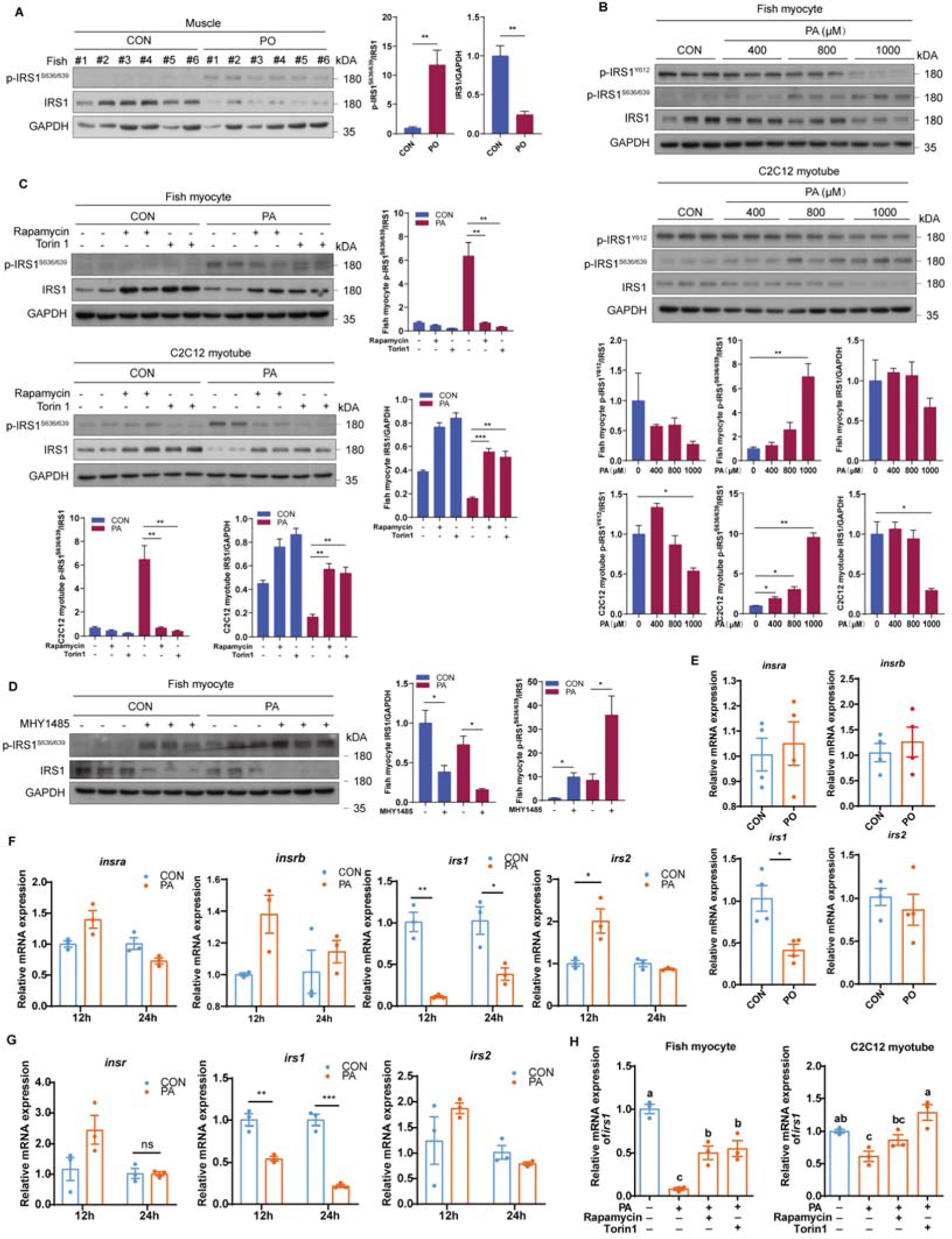
PA-induced insulin resistance is dependent on the negative regulation of IRS1 by mTORC1. (A) Immunoblotting of IRS1 phosphorylation and protein levels in the muscle of fish fed CON or PO diet (n=6). (B) IRS1 phosphorylation and protein levels were assayed by immunoblotting in fish myocytes and C2C12 myotubes treated with the indicated concentrations of PA for 12 h(n=3). (C) IRS1 phosphorylation and protein levels were measured by immunoblotting in fish myocytes and C2C12 myotubes treated with rapamycin or Torin1 treatment in the presence or absence of PA for 12 h (n=3). (D) IRS1 phosphorylation and protein levels were tested by immunoblotting in fish myocytes treated with control or MHY1485 treatment in the presence or absence of PA for 12 h (n=3). (E) Relative mRNA levels of *insa*, *insb*, *irs1* and *irs2* were tested by quantitative PCR in the muscle of fish fed CON or PO diet (n=4). (F) Relative mRNA levels of *insa*, *insb*, *irs1* and *irs2* were analyzed by quantitative PCR in fish myocytes under control or PA treatments for 12 h and 24 h (n=3). (G) Relative mRNA levels of *insr*, *irs1* and *irs2* were measured by quantitative PCR in C2C12 myotubes under control or PA treatments for 12 h and 24 h (n=3). (H) Relative mRNA levels of *irs1* were analyzed by quantitative PCR in fish myocytes and C2C12 myotubes treated with control, rapamycin or Torin1 in the presence or absence of PA for 12 h (n=3). The results are presented as the mean ± SEM and were analyzed using independent *t*-tests (\**p* < 0.05, \*\**p* < 0.01, \*\*\**p* < 0.001) and Tukey’s tests (bars bearing different letters are significantly different among treatments (*p* < 0.05)). See also Figure S6.

### PA inhibits the nuclear translocation of TFEB to impede IRS1 transcription

Given that downstream transcription factors of mTORC1 mediate the regulation of gene transcription, we explored which downstream transcription factor is involved in the inhibition of IRS1 transcription under PA treatment. Dual luciferase experiments in HEK293T cells showed that TFEB had the strongest ability to elevate the luciferase activity of the IRS1 promoter among the crucial downstream transcription factors of mTORC1 (Figure 7A). Moreover, TFEB enhanced the promoter activity of IRS1 in a dose-dependent manner (Figure 7B), and mutations of the predicted TFEB binding site 4 and site 6 in IRS1 promoter significantly reduced the promoter activity of IRS1 in HEK293T cells (Figure 7C). Furthermore, ChIP and EMSA experiments in HEK293T cells verified that TFEB can directly bind to the IRS1 promoter at site 4 and site 6 (Figures 7D and 7E). Importantly, overexpression of TFEB in fish myocytes also strongly increased the protein expression levels of IRS1 (Figure 7F). These results suggest that TFEB can promote IRS1 transcription through binding to the IRS1 promoter region. Next, the effect of PA on TFEB cellular localization was assayed via cell fractionation analyses and the results showed that PA treatment prevented the nuclear translocation of TFEB in fish myocytes and C2C12 myotubes (Figure 7G). Considering that mTORC1 controls TFEB nuclear translocation by phosphorylating TFEB at Ser211^52^, fish myocytes and C2C12 myotubes were treated with rapamycin or Torin1 in the presence of PA. These two inhibitors improved the nuclear translocation of TFEB under PA treatment (Figure 7H), suggesting that PA-induced mTORC1 activation contributes to defective nuclear translocation of TFEB, which may induce suppression of IRS1 transcription and subsequent insulin resistance. To further prove this notion, cultured cells were treated with TFEB activator 1, a synthesized curcumin derivative that can specifically bind to TFEB and induce TFEB nuclear translocation^53^. The results showed that TFEB activator 1 treatment blocked the decline in IRS1 mRNA and protein expression levels under PA condition (Figures 7I and 7J). Furthermore, insulin-stimulated phosphorylation levels of AKT in the presence of PA were attenuated by TFEB activator 1 treatment in fish myocytes and C2C12 myotubes (Figure 7K). Taken together, these results indicate that mTORC1 activation induced by PA leads to cytoplasmic localization of TFEB which inhibits IRS1 transcription and causes insulin resistance.

**Figure 7.**
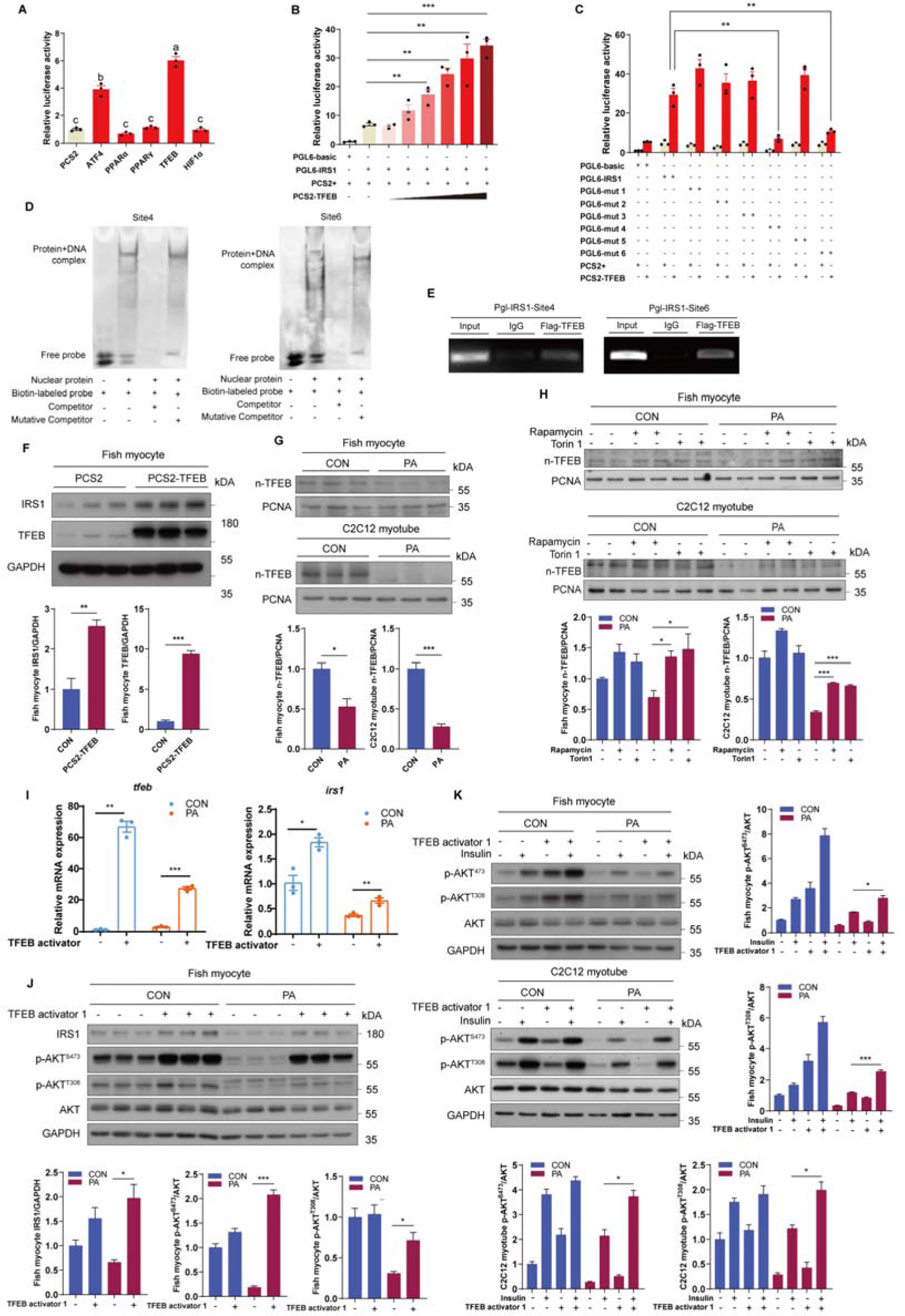
PA inhibits the nuclear translocation of TFEB to impede IRS1 transcription. (A) Relative dual luciferase activity analysis was conducted to measure the effect of ATF4, PPARα, PPARγ, TFEB and HNF1α on IRS1 promoter activity in HEK293T cells (n=3). (B) Relative dual luciferase activity analysis was conducted to measure the effect of TFEB at different concentration gradients on IRS1 promoter activity in HEK293T cells (n = 3). (C) Relative dual luciferase activity analysis was performed to test the effect of TFEB on IRS1 promoter activity with mutation of the predicted binding sites in HEK293T cells (n = 3). (D and E) The binding between TFEB and the predicted region of the IRS1 promoter was demonstrated by EMSA (D) and ChIP (E) in HEK293T cells. (F) TFEB and IRS1 protein levels were measured by immunoblotting in fish myocytes transfected with pCS2 (empty vector) or pCS2-TFEB plasmids (n = 3). (G) TFEB nuclear translocation was assayed by immunoblotting nuclear fractions of fish myocytes and C2C12 myotubes treated with control or PA treatment for 12 h (n = 3). (H) TFEB nuclear translocation was tested by immunoblotting nuclear fractions of fish myocytes and C2C12 myotubes treated with control, rapamycin or Torin1 treatment in the presence or absence of PA for 12 h (n = 3). (I) Relative mRNA levels of *tfeb* and *irs1* were analyzed by quantitative PCR in fish myocytes treated with control or TFEB activator 1 treatment in the presence or absence of PA for 12 h (n = 3). (J) IRS1 protein levels and the phosphorylation of AKT were tested by immunoblotting in fish myocytes treated with control or TFEB activator 1 treatment in the presence or absence of PA for 12 h (n = 3). (K) AKT phosphorylation levels were measured by immunoblotting in fish myocytes and C2C12 myotubes (n = 3). Cells were pretreated with control or TFEB activator 1 treatment in the presence or absence of PA for 12 h, and then stimulated with insulin for 5 min. The results are presented as the mean ± SEM and were analyzed using independent *t*-tests (\**p* < 0.05, \*\**p* < 0.01, \*\*\**p* < 0.001) and Tukey’s tests (bars bearing different letters are significantly different among treatments (*p* < 0.05)).

## Discussion

Dietary habits can affect the metabolic homeostasis and are associated with multiple diseases. Previous studies in mammals have shown that dietary HFD can lead to severe insulin resistance and glucose intolerance^54, 55^, indicating a strong correlation between lipid overload and the development of T2D^13^. However, not all types of fatty acids can induce insulin resistance. Studies in humans have shown that replacing a monounsaturated fatty acid (MFA) diet with a SFA diet can lead to the impairment of insulin pathway, and in human skeletal muscle, dietary PA also induces more extreme insulin resistance than OA^6, 56^. These studies suggest that the divergent effects of fatty acids on insulin signaling may depend on saturation, whereas the underlying mechanisms remain largely obscure. Here, consistent with previous studies, we found that PA (SFA) contributed to systemic and cellular insulin resistance, but OA (MFA) or LA (polyunsaturated fatty acid) had no impact on the insulin sensitivity *in vivo* and *in vitro*. Notably, we also observed that in contrast to other AKT downstream kinases, the activity of mTORC1 was boosted in a time- and dose-dependent manner under PA treatments. However, OA and LA failed to promote mTORC1 activity. Moreover, inhibition of mTORC1 by rapamycin attenuated PA-induced insulin resistance, while activating mTORC1 by MHY1485 aggravated PA-induced insulin resistance. These results indicate that SFA-induced insulin resistance is dependent on the hyperactivation of mTORC1.

mTORC1 is a critical connection between nutritional status and metabolic control. As several amino acid sensors have been identified in recent years, the mechanisms of amino acid-induced mTORC1 activation have been well established^15^. However, progress in determining the mechanisms of fatty acid-induced mTORC1 activation is limited. Only one recent study suggested that PA can induce the activation of mTORC1 through STING1-TBK1-SQSTM1 pathway^57^, and another study indicated that fatty acid-mediated regulation of mTORC1 activity was dependent on the de novo synthesis of phosphatidic acid^58^. But it is still unknown whether there are other mechanisms involved in the activation of mTORC1 by fatty acids. In this study, we found that blocked mitochondrial fatty acid β oxidation inhibited PA-induced mTORC1 activation. Moreover, we also discovered that the reduction of mTORC1 activity by blocking fatty acid β oxidation was rescued through sodium acetate treatment under PA conditions and suppression of ACLY diminished PA-induced mTORC1 activation. Thus, these results indicate that PA-induced mTORC1 activation is dependent on mitochondrial fatty acid β oxidation and that acetyl-CoA plays a crucial role in coupling mitochondrial fatty acid oxidation and mTORC1 activity.

Growing lines of evidence suggested a strong link between mitochondrial fatty acid oxidation and mTORC1 signaling^59^. As a central regulator of anabolism, mTORC1 is considered to inhibit fatty acid β oxidation pathway for energy storage or ketogenesis^60^. Several studies revealed that restrained mTORC1 by rapamycin induced fatty acid β oxidation in rat hepatocytes through increasing expression of fatty acid β oxidation related enzymes^61, 62^. Likewise, mice with whole-body knockout of S6K1 showed enhanced fatty acid β oxidation and increased expression levels of CPT1 in isolated adipocytes^21^, and S6K1/S6K2 double-knockout mice also exhibited elevated fatty acid β oxidation of in isolated myoblasts by activating AMPK^60^. Furthermore, a recent study has established that FOXK1 can mediate the inhibition of fatty acid β oxidation by mTORC1^63^. Thus, these collective data revealed that fatty acid β oxidation was restrained by mTORC1. However, conversely, the role of mitochondrial fatty acid oxidation in the regulation of mTORC1 is still controversy. A study in prostate cancer cells suggested that inhibited fatty acid β oxidation by etomoxir reduced mTORC1 activity^64^, and another study found that deleting CPT1B specifically in skeletal muscle of mice suppressed mTORC1 by provoking AMPK activation^65^. Consistent with these studies, our results showed that acetyl-CoA derived from mitochondrial fatty acid β oxidation induced mTORC1 activation under PA treatment, indicating that acetyl-CoA may be a novel insight linking fatty acid β oxidation and mTORC1 signaling. Paradoxically, unlike other studies, a recent study found that mice with heart specific CPT2-deficient exhibited induction of mTORC1 pathway. Thus, the effects of fatty acid β oxidation on mTORC1 pathway are complicated and may differ under variable physiological and pathological conditions. Further studies are needed to determine the sophisticated mechanisms underlying the regulation of fatty acid β oxidation on mTORC1 signaling.

In this study, we also found that PA, OA or LA had different effects on mitochondrial fatty acid β oxidation. Using LC–MS, we showed that PA treatment increased the contents of short/medium-chain acyl-CoA and acylcarnitine in comparison with OA or LA treatment. Moreover, seahorse real-time cell metabolic analysis showed that PA treatment elevated mitochondrial OCR and maximal oxygen consumption rates compared with OA or LA. Likewise, PA-induced increase of fatty acid oxidation-related gene expressions was more robust than OA or LA *in vivo* and *in vitro*. Thus, these results suggested that although all three fatty acids can be oxidation in mitochondria, PA may be preferred to enter the mitochondria for fatty acid β oxidation, compared with OA or LA. Previous studies have found that OA is more inclined to synthesize triglycerides to induce the formation of lipid droplets than PA^66, 67^. Likewise, we also found that OA significantly increased the contents of 18:1-CoA in comparison with PA. Thus, we speculate that, after entering the cell, OA is more preferentially synthesized to triglyceride for storage than fatty acid oxidation. Moreover, LA is considered to be a precursor of arachidonic acid, and can be converted to a myriad of bioactive compounds called eicosanoids^68^. Similarly, we found that LA markedly elevated the contents of 18:2-CoA/18:3-CoA. Thus, we conjecture that LA preferentially synthesizes functional lipids compared to entering mitochondria for fatty acid oxidation. Together, differences in the levels of acetyl-CoA produced by these three fatty acids may be related to their metabolic pathway preferences. There may be two reasons for why PA prefers to enter mitochondrial for fatty acid oxidation. On one hand, due to differences in the structure of PA, OA and LA, the substrate affinity of CPT1B to these fatty acyl-CoAs may be different, that may contribute to the different rates of fatty acid to enter into mitochondria. On the other hand, in contrast to the β-oxidation of SFAs, the β-oxidation of UFAs requires the involvement of 2,4-dienoyl-CoA reductase^69^, and thus the β-oxidation of SFAs may be more efficient. However, the current understanding of differences in fatty acid oxidation between SFAs and UFAs is insufficient, so more studies are needed in the future to further explore the underling mechanisms behind these differences.

Previous studies have shown that impaired mitochondrial fatty acid oxidation contributes to insulin resistance and that improved mitochondrial fatty acid oxidation capacity can ameliorate insulin resistance provoked by diet or obesity^70, 71^. However, some studies challenged this theory by showing that CPT1B or CPT2 muscle-specific knockout mice exhibited resistance to diet-induced insulin resistance^72, 73^. Moreover, mice lacking malonyl-CoA decarboxylase (MCD), an enzyme that stimulates fatty acid oxidation by reducing malonyl-CoA-mediated restriction of CPT1, exhibited improved glucose intolerance provoked by diet^74^. In the present study, we observed that restraining fatty acid β oxidation enhanced insulin-stimulated glucose uptake and phosphorylation of AKT under PA conditions. Furthermore, sodium acetate addition blocked the recovery of insulin-stimulated glucose uptake and phosphorylation of AKT by perhexiline maleate. Thus, our results indicate that PA-induced insulin resistance is dependent on mitochondrial fatty acid β oxidation and that targeting fatty acid β oxidation may be a potential therapeutic strategy for SFA diet-induced insulin resistance. These results also further reveal a novel role for acetyl-CoA in mediating the link between fatty acid β oxidation and insulin resistance.

Acetyl-CoA is not only a metabolite of the TCA cycle, but also serves as a substrate for acetylation modification. Several studies have revealed that acetyl-CoA derived from fatty acid oxidation can regulate cellular functions under physiological or pathological conditions by altering protein acetylation modifications. For example, acetyl-CoA derived from fatty acid oxidation promoted lymphangiogenesis by facilitating the acetylation of histones by histone acetyltransferase p300 at lymphangiogenic genes^75^, and acetyl-CoA produced by fatty acid oxidation contributed to aggressive growth of glioblastoma multiforme by upregulating NF-κB/RelA acetylation^76^. Our previous study also showed that acetyl-CoA derived from fatty acid oxidation increased p65 acetylation to intensify inflammation^77^. In addition, acetylation modification can also regulate mTORC1 activity through acetylating mTORC1 components or their associated proteins including Raptor and Rheb^32, 35^. Therefore, we hypothesized that acetylation modification may mediate PA-induced mTORC1 activity. The results showed that PA enhanced the acetylation levels of Rheb in a dose-dependent manner but had no effect on Raptor acetylation. Likewise, restriction of fatty acid oxidation attenuated the induction of Rheb acetylation under PA treatment and sodium acetate treatment aggravated Rheb acetylation in the presence or absence of PA treatment. As previous studies have shown that Rheb acetylation is essential for mTORC1 activation and that Rheb is also a highly conserved protein^35^, we consider that acetyl-CoA derived from fatty acid oxidation upregulates Rheb acetylation which induces mTORC1 activation under PA condition.

In addition to the substrate acetyl-CoA, acetylation modification usually requires the engagement of acetyltransferases. A recent study showed that CBP has the strongest ability to acetylate Rheb in HEK293T cells^35^. However, in the present study, inhibition of CBP by C646 and spermidine failed to alleviate PA-induced mTORC1 activation, suggesting that other acetyltransferases may mediate Rheb acetylation under PA condition. Subsequent results found that suppression of Tip60 by pharmacological inhibitors or siRNA knockdown relieved the induction of mTORC1 under PA treatment. Consistently, suppression of Tip60 by MG149 treatment or siRNA knockdown reduced PA-induced Rheb acetylation. Furthermore, Co-IP assays also demonstrated the interaction between Tip60 and Rheb. Taken together, these results indicate that Tip60 mediates Rheb acetylation to activate mTORC1 signaling under PA condition. In line with our results, a recent study also found that Tip60 can regulate triacylglycerol synthesis by acetylating lipin 1 in response to fatty acids stimulation^78^. Therefore, Tip60 may be a vital node connecting fatty acid sensing and acetylation modification. In this study, we found that acetyl-CoA derived from mitochondrial fatty acid β oxidation fueled Tip60-mediated Rheb acetylation to induce mTORC1 activation under PA treatment, which may provide novel insight into the mechanism of lipid sensing by mTORC1. Furthermore, we also found that reducing Rheb acetylation by inhibiting ACLY or Tip60 ameliorated PA-induced insulin resistance. Considering that chronic suppression of mTORC1 by pharmacological inhibitors such as rapamycin leads to glucose intolerance through hindering mTORC2, targeting mitochondrial fatty acid β oxidation mediated acetyl-CoA production or Tip60-mediated Rheb acetylation may provide novel therapeutic opportunities for lipid surplus induced T2D.

Aberrant regulation of mTORC1 is associated with the development of T2D. Numerous studies have unveiled that mTORC1 activation induces phosphorylation-dependent degradation of IRS1 to impede insulin signaling^20, 21^. However, whether mTORC1 signaling regulates the transcription of IRS1 remains elusive. Consistent with these studies, our current results also showed that PA treatment reinforced the S636/S639 phosphorylation of IRS1 in an mTORC1-dependent manner. Notably, the mRNA expression of *irs1* was decreased by PA and rapamycin or Torin1 treatment upregulated the mRNA expression of *irs1* under PA treatment, indicating that mTORC1 signaling also mediated the transcriptional regulation of IRS1. Furthermore, we identified TFEB may play an important role in IRS1 transcription and this notion was strongly supported by the results from ChIP and EMSA assays. Moreover, current study also showed that PA treatment inhibited the nuclear translocation of TFEB by activating mTORC1 pathway and that enhancing TFEB expression by TFEB activator 1 attenuated PA-induced glucose intolerance and insulin resistance. Consistent with our results, a study in adipose tissue macrophages showed that lysosomal stress response provokes TFEB-GDF15 to protect against obesity and insulin resistance^79^. Therefore, besides regulating lysosomal biogenesis and autophagy, TFEB also serves as a vital modulator of glucose homeostasis and insulin sensitivity, which may provide a novel mechanistic clue for developing therapeutic strategies to combat T2D.

In summary, our work unveils an evolutionarily conserved mechanism by which mitochondrial fatty acid β oxidation flux of acetyl-CoA induces mTORC1 activation through enhancing Tip60-mediated Rheb acetylation under PA condition. Subsequently, hyperactivation of mTORC1 boosted serine phosphorylation of IRS1 and inhibited TFEB-mediated transcription of IRS1, leading to insulin resistance (Figure 8). These findings may provide insight for understanding the mechanism of SFA- or lipid surplus-induced insulin resistance and open a promising therapeutic avenue to improve insulin sensitivity and glucose tolerance.

**Figure 8.**
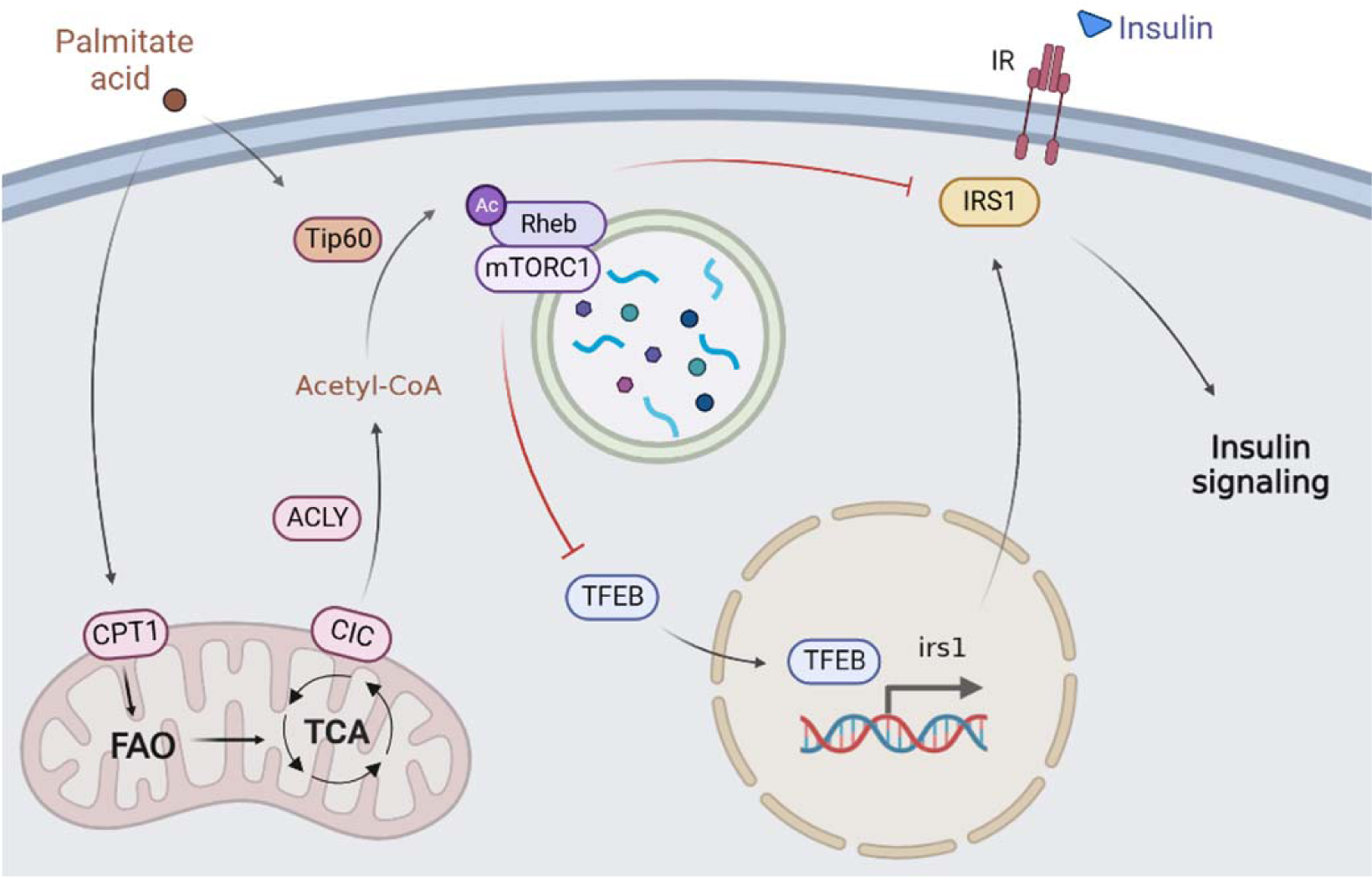
A working model of how excessive PA induces insulin resistance.

## Methods

### Animal studies

Four-month-old large yellow croakers of similar size (mean weight 15.67 ± 0.11 g) were obtained from the Aquatic Seeds Farm of the Marine and Fishery Science and Technology Innovation Base (Ningbo, Zhejiang, China) and bred in floating sea cages at 26.8 ± 3°C, 30.8-35.7‰ salinity and 6–7 mg/L dissolved oxygen. The fish were randomly divided into four groups that were fed diverse experimental diets. Four isonitrogenous (42% crude protein) and isolipidic (12% crude lipid) experimental diets were formulated, containing control diet (fish oil as a source of dietary fat) which is suitable for large yellow croaker growth, PA rich diet (palm oil as a source of dietary fat), OA rich diet (olive oil as a source of dietary fat) and LA rich diet (soybean oil as a source of dietary fat). The fatty acid profiles of these four diets are listed in Table S1. The male and female fish were fed each diet twice a day at 05:00 and 17:00 for 10 weeks. At the end of the feeding trial, MS222 (1:10, 000; Sigma, USA) was used to anesthetize the fish, and the liver, muscle and plasma of these fish were sampled for subsequent analysis.

DsRNA was synthesized using the TranscriptAid T7 High Yield Transcription Kit (Thermo Fisher Scientific, USA) according to the manufacturer’s instructions. Fish were intraperitoneally injected with dsRNA-control, dsRNA-CPT1B or dsRNA-ACLY for 36 h at a dose of 2 μg/g according to the body weight. Sampling collection was the same as described above.

In the current study, all experimental procedures performed on fish were conducted in strict accordance with the Management Rule of Laboratory Animals (Chinese Order No. 676 of the State Council, revised 1 March 2017).

### Cell culture

Primary myocytes were isolated from large yellow croaker according to the following methods. Muscle tissues were removed and placed in sterile phosphate buffer (PBS, Biological Industries, Israel) containing penicillin (Solarbio, China) and streptomycin (Solarbio, China). Then, tissues were cut into small pieces in Dulbecco’s modified Eagle medium/Ham’s F12 medium (1:1) (DMEM/F12, Biological Industries) and washed twice with DMEM/F12 medium to remove serum. Subsequently, tissues were digested with 0.2% trypsin (Thermo Fisher Scientific, USA) for 20 min and washed twice with DMEM/F12 medium. Later, after digestion with 0.1% trypsin for another 10 min and neutralization with DMEM/F12 medium with fetal bovine serum (FBS, Biological Industries), the cell precipitates were resuspended in DMEM complete medium composed of DMEM/F12 medium supplemented with 15% FBS, 100 U penicillin and 100 mg/mL streptomycin. The cell suspension was inoculated into a six-well culture plate and incubated at 28°C under 5% CO_2_.

Mouse C2C12 myoblast cells were obtained from the Cell Bank of the Chinese Academy of Sciences (Shanghai, China) and were cultured in Dulbecco’s modified Eagle medium (DMEM, Biological Industries) supplemented with 10% fetal bovine serum, 100 units/mL penicillin and 100 mg/mL streptomycin within an atmosphere of 5% CO_2_ at 37°C. To induce differentiation and myotube formation, 10% fetal bovine serum was substituted by 2% horse serum (Gibco, USA) in DMEM with penicillin and streptomycin. After 5 days, the differentiated myotubes were used for subsequent assays.

HEK293T cells were obtained from the Cell Bank of the Chinese Academy of Sciences (Shanghai, China) and were cultured in DMEM supplemented with 10% FBS, 100 units/mL penicillin and 100 mg/mL streptomycin at 37°C with 5% CO_2_.

### Cell treatments

For PA, OA or LA *in vitro* treatment, fatty acid free BSA (Equitech-Bio, USA) was dissolved in FBS-free DMEM at room temperature according the ratio 1:100 (1 g fatty-acid free BSA: 100 ml FBS-free DMEM). 500 mg PA (Merck, Cat#P0500), OA (Merck, Cat#O1008) or LA (Merck, Cat#L1376) was dissolved in 10 ml ethanol to obtain PA, OA or LA stock solution respectively. Then PA, OA or LA stock solution was blow-drying with nitrogen gas and was dissolved in 0.1 M NaOH and warming at 75°C until clear to obtain 100 mM PA, OA or LA solution. Subsequently, 100 mM PA, OA or LA solution was added to 1% BSA solution according the ratio 1:100 (100 mM PA:1% BSA, v/v) at 50°C. Finally, the mixture was filtered using a 0.45 μM filter and stored at −20°C. Fatty acid treatment was carried out by incubating fish primary myocytes or C2C12 myotubes with serum free media containing the indicated concentrations of PA, OA or LA for 12 h-24 h.

For insulin *in vitro* treatment, insulin powder (Merck, USA) was dissolved in hydrochloric acid (pH=2) to obtain 1 mg/ml stock solution. Insulin stimulation was performed by treating fish primary myocytes or C2C12 myotubes with 100 nM insulin for 5 min.

For rapamycin or Torin1 *in vitro* treatment, rapamycin (Med Chem Express, #HY-10219, USA) or Torin1 (Med Chem Express, #HY-13003, USA) was dissolved in dimethyl sulfoxide (DMSO, Solarbio, China) to obtain 1 mM stock solution respectively. To inhibit mTORC1, 500 nM rapamycin or 500 nM Torin1 treatment was added to the culture medium of fish primary myocytes or C2C12 myotubes for 12 h in the presence or absence of PA.

For MHY1485 *in vitro* treatment, MHY1485 (Med Chem Express, #HY-B0795, USA) was dissolved in DMSO (Solarbio, China) to obtain 10 mM stock solutions. To activate mTORC1, 10 μM MHY1485 was added to the culture medium of fish primary myocyte for 12 h in the presence or absence of PA.

For etomoxir or perhexiline maleate *in vitro* treatments, etomoxir (Med Chem Express, #HY-50202, USA) or perhexiline maleate (Med Chem Express, #HY-B1334A, USA) was dissolved in DMSO (Solarbio, China) to obtain 50 mM stock solution respectively. To inhibit mitochondrial fatty acid β oxidation, 50 μM etomoxir or 25 μM perhexiline maleate was added to the culture medium of fish primary myocytes or C2C12 myotubes for 12 h in the presence or absence of PA.

For BMS-303141 treatment, BMS-303141 (Med Chem Express, #HY-16107, USA) was dissolved in DMSO (Solarbio, China) to obtain 25 mM stock solutions. To inhibit ACLY, 25 μM BMS-303141 was added to the culture medium of fish primary myocytes for 12 h in the presence or absence of PA.

For sodium acetate treatment, sodium acetate (Merck, #S2889, USA) was dissolved in ultrapure water from a Milli-Q water system to obtain 5M stock solution. To increase the content of cellular acetyl-CoA, the indicated concentrations of sodium acetate were added to the culture medium of fish primary myocytes and C2C12 myotubes for 12 h in the presence or absence of PA.

For C646, spermidine or MB-3 treatment, C646 (Med Chem Express, #HY-13823, USA), spermidine (Med Chem Express, #HY-B1776, USA) or MB-3 (Merck, #M2449, USA) was dissolved in DMSO (Solarbio, China) to obtain 50 mM stock solution respectively. To inhibit CBP/P300, the indicated concentrations of C646 or spermidine were added to the culture medium of C2C12 myotubes for 12 h in the presence of PA. To inhibit GCN5, the indicated concentrations of MB-3 were added to the culture medium of C2C12 myotubes for 12 h in the presence of PA.

For MG149 treatment, MG149 (Med Chem Express, #HY-15887, USA) was dissolved in DMSO (Solarbio, China) to obtain 150 mM stock solution. To inhibit Tip60, 150 μM MG149 was added to the culture medium of fish primary myocytes or C2C12 myotubes for 12 h in the presence or absence of PA.

For TFEB activator 1 treatment, TFEB activator 1 (Med Chem Express, #HY-135825) was dissolved in DMSO (Solarbio, China) to obtain 10 mM stock solution. To activate TFEB, 15 μM TFEB activator 1 was added to the culture medium of fish primary myocytes and C2C12 myotubes for 12 h in the presence or absence of PA.

### siRNA transfection

C2C12 cells were seeded in plates with DMEM containing 10% FBS and then differentiated for 5 days with DMEM containing 2% horse serum. Differentiated myotubes were transiently transfected with siRNAs (siRNAs against CPT1B, CPT2, ACLY or Tip60 and scramble siRNA were commercially synthesized (GenePharma, China)) using Lipofectamine™ RNAiMAX (Invitrogen, USA) according to the manufacturer’s instructions. Knockdown efficiency was verified by quantitative PCR and immunoblotting. The siRNA sequences used are listed in Table S2.

### NEFA, TG, glucose and insulin content assays

The plasma NEFA level was measured by Non-esterified Fatty Acids (NEFA) Assay Kit according to the manufacturer’s instructions (Nanjing Jiancheng Bio-Engineering Institute, China). The TG content was analyzed by Triglyceride (TG) Content Assay Kit (Applygen Technologies Inc., China). The fasted blood glucose and insulin contents were tested by Glucose Assay Kit (Applygen Technologies Inc., China) and Fish INS ELISA Assay Kit (CUSABIO Technology, USA) according to the manufacturer’s instructions.

### ITT and GTT

Assays were carried out on large yellow croaker fed CON diet or PO diet after 10 weeks. For GTTs, fish were fasted for 24 h and then intraperitoneally injected with glucose (0.9g/kg body weight). Blood was sampled at 0 h, 0.5 h, 1 h, 2 h, 4 h and 8 h after injection with glucose. For ITTs, fish were fasted for 24 h and then intraperitoneally injected with insulin (0.052mg/kg body weight). Blood was sampled at 0 h, 0.5 h, 1 h, 2 h, 4 h and 8 h after injection with insulin.

### Glucose uptake

Glucose uptake was detected by 2-DG uptake assays using a Glucose Uptake-Glo™ Assay (Promega, J1341, USA) according to the manufacturer’s instructions. In brief, cells plated in 96-well plates were incubated with the indicated treatments without serum or glucose and stimulated with 100 nM insulin for 1 h. Then after treatment with 2-DG for 10 min, cells were lysed in stop buffer and neutralized with neutralization buffer. The lysates were treated with 2DG6P detection reagent and luminescence was recorded in 0.5 s intervals.

### Acyl-CoA and acyl-carnitine quantification by LC/MS

Fish myocytes were incubated with 800 μM PA, OA or LA for 12 h before collection and freezing. Analysis of acyl-CoAs and acyl-carnitines was carried out at LipidALL Technologies as previously described^80^. Briefly, 300 µL of extraction buffer containing isopropanol, 50 mM KH2PO4, 50 mg/mL BSA (25:25:1 v/v/v) acidified with glacial acetic acid was added to cells. Next, 19:0-CoA and d3-16:0-carnitine was added as internal standards and lipids were extracted by incubation at 4 °C for 1 h at 1500 rpm. Following this, 300 µL of petroleum ether was added and the sample was centrifuged at 12000 rpm for 2 min at 4 °C. The upper phase was removed. The samples were extracted two more times with petroleum ether as described above. To the lower phase finally remaining, 5 µL of saturated ammonium sulfate was added followed by 600 µL of chloroform:methanol (1:2 v/v). The sample was then incubated on a thermomixer at 450 rpm for 20 min at 25 °C, followed by centrifugation at 12000 rpm for 5 min at 4 °C. Clean supernatant containing long-chain acyl-CoAs was transferred to fresh tube and subsequently dried in the SpeedVac under OH mode (Genevac). To improve recovery of polar short-chain CoAs, the remaining pellet was extracted with 1 ml trichloroacetic acid. The acidic extract was purified by solid phase extraction using Oasis HLB 1cm3 (30 mg) SPE columns from Waters. The purified extract containing polar CoAs was dried in a SpeedVac under OH mode. The two extracts were combined and resuspended in methanol:water (9:1 v/v) containing 0.05% acetic acid, and analyzed on a Shimadzu 40X3B-UPLC coupled to Sciex QTRAP 6500 Plus.

### Metabolic flux analysis

For metabolic-tracing analyses, fish myocytes were exposed to 800 μM [U-^13^C_16_]-palmitate (Merck, #605573, USA) for 24 h. Analysis of acyl-CoAs was carried out at LipidALL Technologies as previously described^80^ and follow the same steps as described in the acyl-CoA and acyl-carnitine quantification by LC/MS.

### Oxygen consumption rate (OCR) measurement

Oxygen consumption rate (OCR) was measured using the seahorse XF 24 Flux Analyzer (Seahorse Biosciences, USA) according to the manufacturer’s protocol. Briefly, fish myocytes were planted in a XF24-well plate (Seahorse Biosciences, USA) and were treated with PA, OA or LA for 12 h before OCR measurement. Then myocytes were incubated with unbuffered assay media at 37°C in ambient CO2 for 1 h. The OCRs of different states were measured by treating myocytes with oligomycin (1.5 μM), FCCP (2 μM) and antimycin & rotenone (0.5 μM), respectively. Myocytes were then collected for determinations of protein content (BCA).

### RNA extraction and reverse transcriptase-quantitative PCR (RT-qPCR)

Total RNA from tissues and cells was extracted using TRIzol reagent (Takara, Japan) and were reversed transcribed into first-strand cDNA using the PrimeScript RT Reagent Kit (Takara) according to the manufacturer’s instructions. RT-qPCR was carried out using SYBR qPCR MasterMix (Vazyme, China). To calculate the expression of genes, the mRNA expression of genes was normalized to the expression of the β-actin gene and the comparative cycle threshold (CT) method (2−▵▵CT method) was employed. The primers used for qPCR are listed in Table S2.

### Acetyl-CoA measurement

The intracellular acetyl-CoA content was assayed using an Acetyl-Coenzyme A Assay Kit (MAK039, Sigma) according to the manufacturer’s instructions. Briefly, the samples were deproteinized by perchloric acid and the Acetyl-CoA Quencher, and Quench Remover were added to the samples to correct for background. Then the samples were mixed with reaction buffer and incubated for 10 min at 37°C. The fluorescence was tested using a plate reader and the following settings: λex535 nm; λem 587 nm.

### Western blot analysis

Total proteins were extracted from tissues and cells using RIPA lysis buffer (Solarbio, China) containing protease and phosphatase inhibitors (Roche, Germany). Nuclear protein was collected using NE-PER Nuclear and Cytoplasmic Extraction Reagents (Thermo Fisher Scientific, USA) according to the manufacturer’s instructions. Equivalent amounts of denatured protein homogenate were resolved by SDS-PAGE on 10% polyacrylamide gels and transferred to a polyvinylidene difluoride membrane (Millipore, USA). After blocking at room temperature for 2 h, the membranes were incubated with primary antibodies overnight. Then the membranes were incubated with secondary antibodies and visualized with BeyoECL Plus Reagent (Beyotime Biotechnology, China). Antibodies against Phospho-AKT (Ser473) (#4060), Phospho-AKT (Thr308) (#13038), AKT (#9272), Phospho-S6K (Thr389) (#9205), S6K (#9202), Phospho-S6 (Ser240/244) (#5364), S6 (#2217), Phospho-GSK-3β (Ser9) (#5558), GSK-3β (#12456), Phospho-AS160 (Thr642) (#8881), AS160 (#2670), Rheb (#13879), Raptor (#2280), Phospho-IRS-1 (Ser636/639) (#2388), IRS1 (#3407), PCNA (#13110), Acetylated-Lysine (#9441), DYKDDDDK Tag (#14793) and HA Tag (#3724) were purchased from Cell Signaling Technology (USA). Antibodies against CPT1B (#22170-1-AP), CPT2 (#26555-1-AP) and Tip60 (#10827-1-AP) were purchased from Proteintech (USA). Antibodies against Phospho-IRS1 (Y612) (#MAB7314) was purchased from RnD systems (USA). Antibody against TFEB (#NB100-1030) was obtained from Novus (USA). Antibody against GAPDH (#TA-08) and HRP-conjugated secondary antibodies were purchased from Golden Bridge Biotechnology (China).

### Plasmid constructs

For expression plasmids, the open reading frames (ORFs) of ATF4, PPARα, PPARγ, TFEB and HIF1α of the large yellow croaker were amplified and subcloned into the PCS2 vector. A FLAG tag was inserted into the pcDN13.1-TFEB expression plasmid. The pcDNA3.1-Tip60 (mouse)-HA plasmid was purchased from Youbio (China) and the pEnCMV-RHEB (mouse)-3×FLAG plasmid was purchased from Miaolingbio (China). For reporter plasmids, the IRS1 wild-type promoter fragment was cloned from the large yellow croaker genomic DNA and then subcloned into the PGL6 vector. The TFEB binding sites on the IRS1 promoter fragment were predicted using the online JASPAR (http://jaspar.genereg.net/) and the IRS1 mutated-type promoter fragments (PGL6-1-mut, PGL6-2-mut, PGL6-3-mut, PGL6-4-mut, PGL6-5-mut, PGL6-6-mut) were generated by *in vitro* site-directed mutagenesis (Vazyme, China). The primers used are listed in Table S2.

### Immunoprecipitation (IP) and co-immunoprecipitation (Co-IP)

For IP analyses, after treatment, cells were lysed with Cell Lysis Buffer for Western Blotting and IP (Beyotime Biotechnology, China) for 20 min. Then, moderate amounts of anti-ac-K antibody agarose beads (Cytoskeleton, Inc., USA) were added to the lysate and incubated for 12 h at 4°C. The immunoprecipitates were washed five times with lysis buffer and mixed with loading buffer. Then, the denatured mixture was analyzed by immunoblotting. For Co-IP, HEK293T cells were lysed with Cell Lysis Buffer for Western Blotting and IP (Beyotime Biotechnology, China) for 20 min after transfection with Rheb-Flag and Tip60-HA for 48 h. Then, the lysate was incubated with ANTI-Flag M2 Affinity Gel (Sigma, USA) or Pierce anti-HA agarose (Thermo Fisher Scientific, USA) at 4°C for 4 h. After washing with the lysis buffer and TBST five times, the binding components were eluted using the Flag peptide (MedChem Express, USA) or HA peptide (MedChem Express, USA) and analyzed by immunoblotting.

### Dual-luciferase reporter assay

HEK293T cells were seeded in 24-well plates and transfected with the expression vector, the promoter reporter vector and the pRL-CMV Renilla luciferase plasmid using Lipofectamine 2000 reagent (Invitrogen, USA). After transfection for 24 h, cells were harvested and the luciferase activity was measured using the Dual-Luciferase Reporter Assay SystemKit (TransGen Biotech Co., Ltd., China) according to the manufacturer’s instructions.

### Chromatin immunoprecipitation assay (ChIP)

The pcDNA3.1-TFEB-Flag vector and PGL6-IRS1 promoter vector were co-transfected into HEK293T cells. After 48 h, the HEK293 cells were fixed with formaldehyde at 37°C for 10 min and were analyzed using a ChIP Assay kit (Beyotime Biotechnology, China) according to the manufacturer’s instructions. Immunoprecipitated DNA was assayed using primers specific for the IRS1 promoter region by PCR. The primers used for ChIP are listed in Table S2.

### Electrophoretic mobility shift assay (EMSA)

HEK293T cells were transfected with the PCS2-TFEB vector. After 48 h, the nuclear protein was collected with NE-PER Nuclear and Cytoplasmic Extraction Reagents (Thermo Fisher Scientific, USA). The sequences of 5′-biotin-labeled double-stranded oligomers are listed in Table S2. Then, the DNA-protein interaction was detected with a LightShiftTM Chemiluminescent electrophoretic mobility shift assay (EMSA) kit (Thermo Fisher Scientific, USA). The primers used for EMSA are listed in Table S2.

### Statistical analysis

The data are presented as the means ± SEM and were analyzed using independent *t*-tests for two groups and one-way ANOVA with Tukey’s test for multiple groups in SPSS 23.0 software. *p*< 0.05 was considered to indicate statistical significance. The number of replicates for each experiment is indicated in the figure legends.

## Acknowledgments

This work was supported by the Key Program of National Natural Science Foundation of China (31830103), National Science Fund for Distinguished Young Scholars of China (31525024), Ten-thousand Talents Program (2018-29), the Agriculture Research System of China (CARS47-11) and Scientific and Technological Innovation of Blue Granary (2018YFD0900402). We also acknowledge Patrick J. Stover (Texas A&M University), Shihuan Kuang (Purdue University), Xiaowei Chen (Peking University), Zhaocai Zhou (Fudan University), Li Xu (Tsinghua University) and Baowei Jiao (Kunming Institute of Zoology, Chinese Academy of Sciences) for their constructive suggestions on the experimental design and revising article. We thank Yanjiao Zhang, Jianlong Du, Yongnan Li, Xiang Xu, Shangzhe Han and Wencong Lai for their experimental assistance.

## Author Contributions

Z.Q.Z. designed the experiments, performed the main experiments and wrote the manuscript. Q.C., X.J.X., W.F., K.C., B.L.L., Q.D.L., Y.T.L. and Y.N.S. conducted other experiments. W.W.D. designed the experiments and revised the manuscript. Y.R.L., W.X. and K.S.M. revised the manuscript. Q.H.A. designed the experiments and wrote the manuscript.

## Supplementary Information File

**Figure S1.**
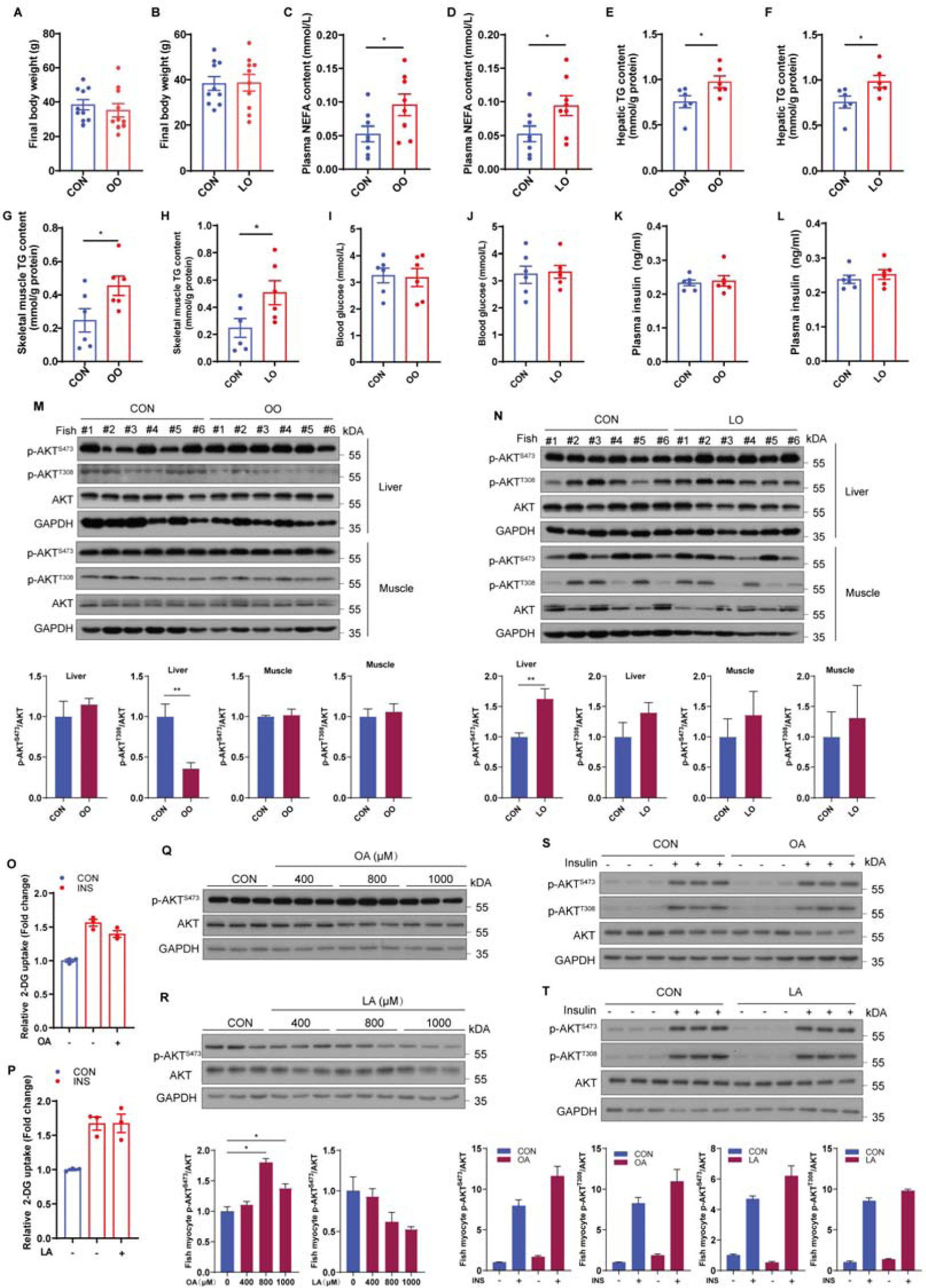
OA and LA have no effect on systemic and cellular glucose homeostasis and insulin sensitivity. Related to Figure 1. (A-N) Fish were fed control (CON), oleic acid (OA) rich (OO) or linoleic acid (LA) rich (LO) diet for 10 weeks. After 12 h fasting, final body weight and blood glucose were measured; plasma, liver and muscle samples were collected. (A and B) Final body weight of fish fed CON, OO or LO diet for 10 weeks (n=10). (C and D) Plasma nonesterified free fatty acid (NEFA) of fish fed different diets (n=8). (E-H) TG levels in liver (E and F) and skeletal muscle (G and H) were tested in fish after treatment with different diets (n=6). (I-L) Blood glucose (I and J) and plasma insulin levels (K and L) were detected in fasted fish fed different diets (n=6). (M-N) Phosphorylation levels of AKT in the liver and skeletal muscle of fish fed different diets were measured by immunoblotting (n=6). (O and P) Insulin-stimulated glucose uptake of fish myocytes was measured by 2-DG uptake assays under control, OA or LA treatment for 12 h (n=3). (Q and R) Phosphorylation levels of AKT in fish myocytes were evaluated by immunoblotting in the presence of the indicated concentrations of OA or LA for 12 h (n=3). (S and T) Phosphorylation levels of AKT in fish myocytes were detected by immunoblotting (n=3). Cells were pretreated with control, OA or LA for 12 h, and then stimulated with insulin for 5 min. The results are presented as the mean ± SEM and were analyzed using independent *t*-tests (\**p* < 0.05; \*\**p* < 0.01).

**Figure S2.**
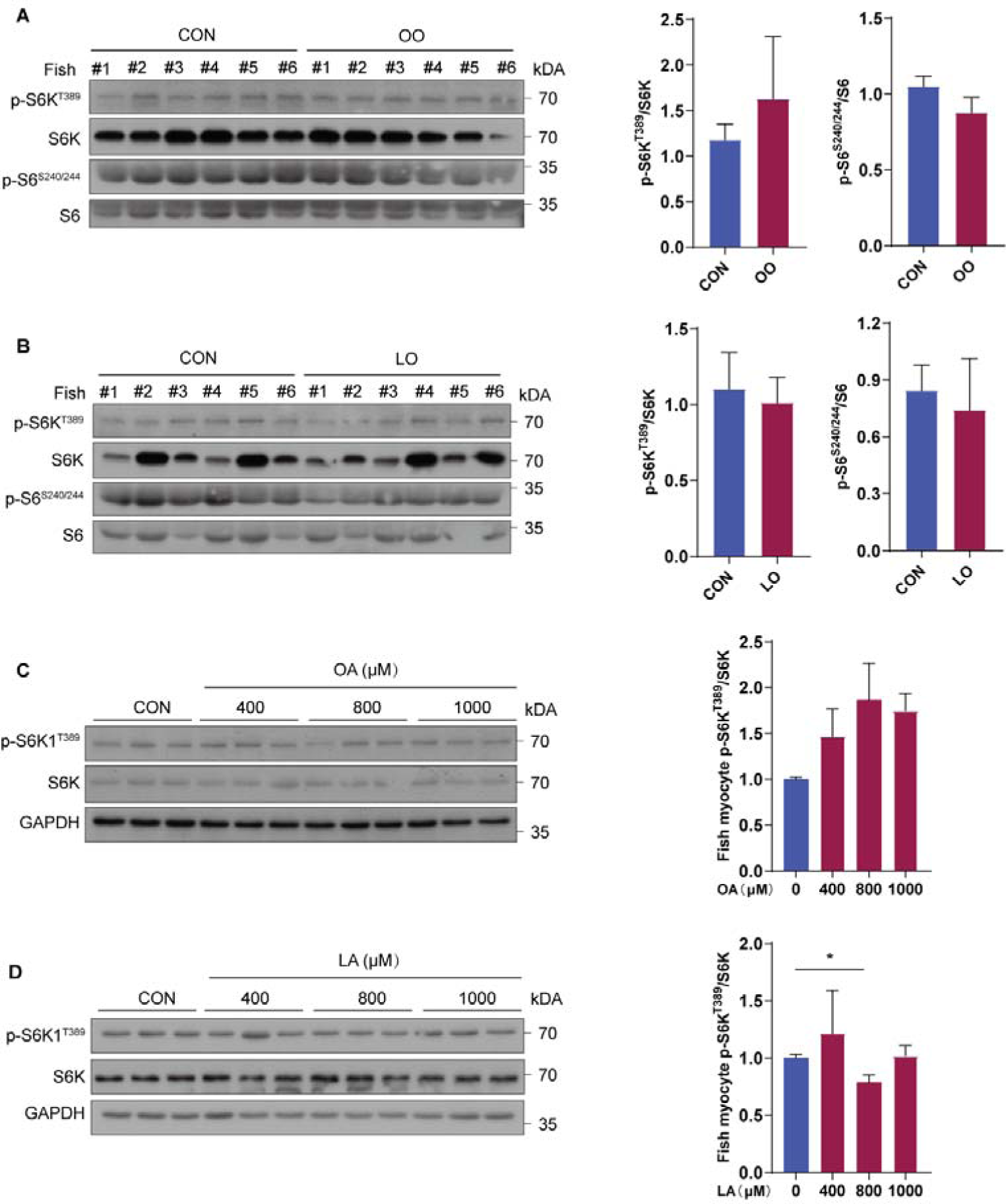
OA and LA have no effect on mTORC1 activity. Related to Figure 2. (A and B) mTORC1 pathway activity was measured by immunoblotting for the phosphorylation of S6K and S6 in skeletal muscle of fish fed CON, OO or LO diet for 10 weeks (n=6). (C and D) mTORC1 pathway activity was analyzed by immunoblotting in fish myocytes treated with the indicated concentrations of OA or LA for 12 h (n=3). The results are presented as the mean ± SEM and were analyzed using independent *t*-tests (\**p* < 0.05).

**Figure S3.**
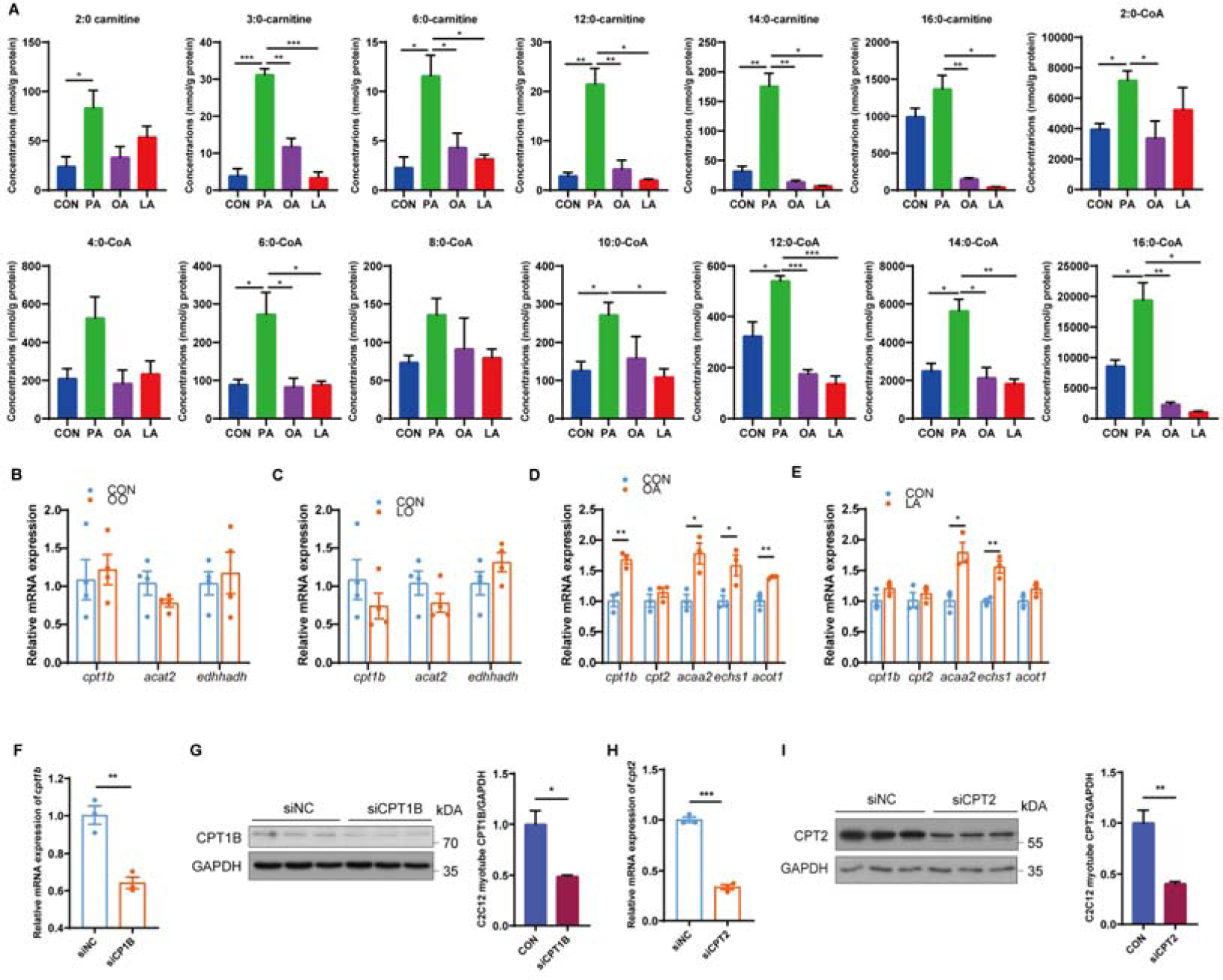
Mitochondrial fatty acid β oxidation is required for PA-induced mTORC1 activation and insulin resistance. Related to Figure 3. (A) The levels of acyl-CoA and acylcarnitine in fish myocytes treated with PA, OA or LA for 12h (n=3). (B and C) Relative mRNA levels of mitochondrial fatty acid β oxidation-related genes were measured by quantitative PCR in the muscle of fish fed CON, OO or LO diet (n=4). (D and E) Relative mRNA levels of mitochondrial fatty acid β oxidation-related genes were examined by quantitative PCR in C2C12 myotubes with control, OA or LA treatment for 12 h (n=3). (F and G) Relative mRNA (A) and protein (B) levels of CPT1B were analyzed by quantitative PCR and immunoblotting in C2C12 myotubes transfected with control siRNA or siRNA against CPT1B for 48 h (n=3). (H and I) Relative mRNA (H) and protein (I) levels of CPT2 were tested by quantitative PCR and immunoblotting in C2C12 myotubes transfected with control siRNA or siRNA against CPT2 for 48 h (n=3). The results are presented as the mean ± SEM and were analyzed using independent *t*-tests (\**p* < 0.05, \*\**p* < 0.01, \*\*\**p* < 0.001).

**Figure S4.**
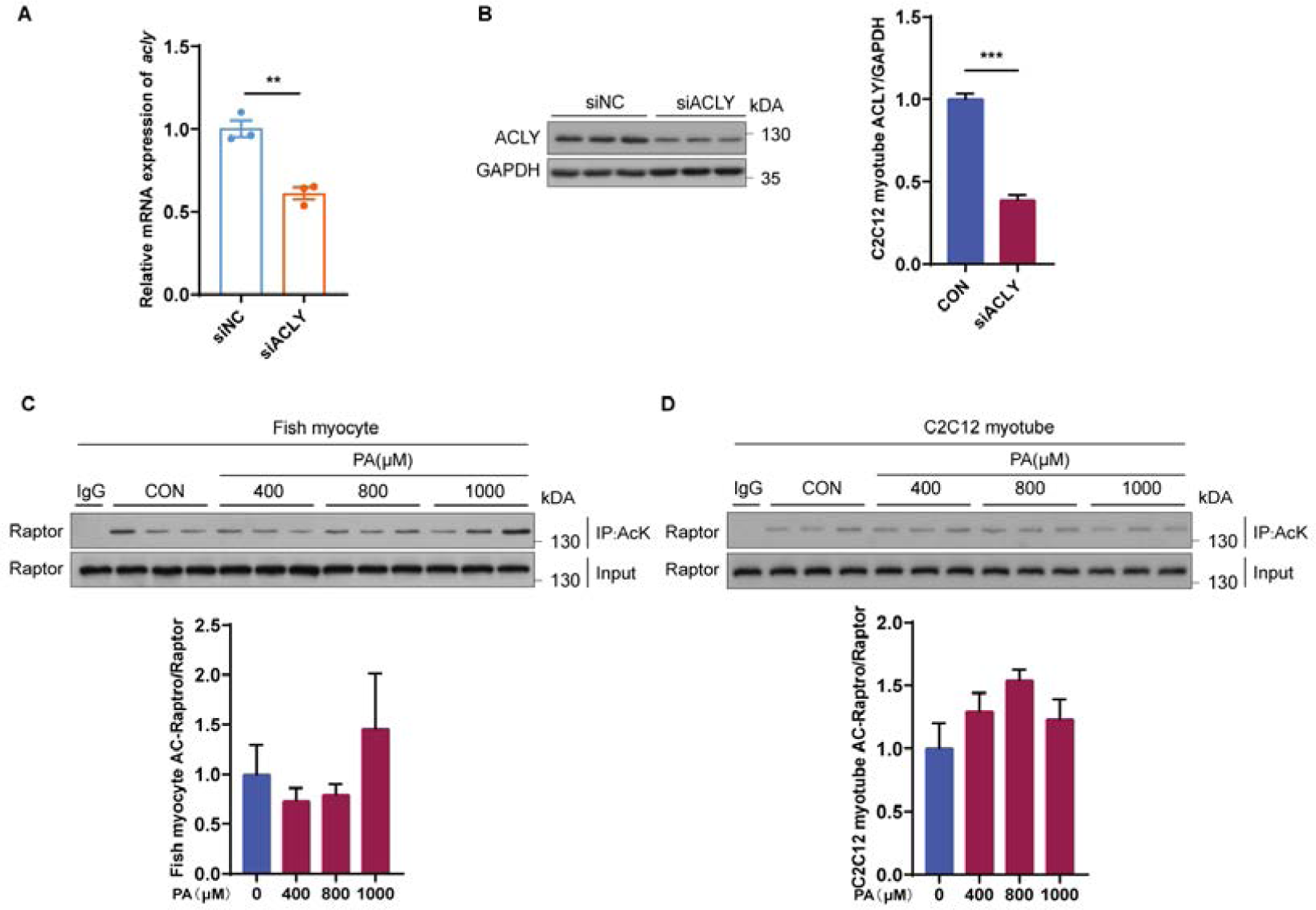
Acetyl-CoA derived from mitochondrial fatty acid β oxidation triggers mTORC1 activation and insulin resistance through enhancing Rheb acetylation. Related to Figure 4. (A and B) Relative mRNA (A) and protein (B) levels of ACLY were analyzed by quantitative PCR and immunoblotting in C2C12 myotubes transfected with control siRNA or siRNA against ACLY for 48 h (n=3). (C and D) Immunoblotting of Raptor acetylation in fish myocytes (C) and C2C12 myotubes (D) with the indicated concentrations of PA for 12 h (n=3). The results are presented as the mean ± SEM and were analyzed using independent *t*-tests (\*\**p* < 0.01, \*\*\**p* < 0.001).

**Figure S5.**
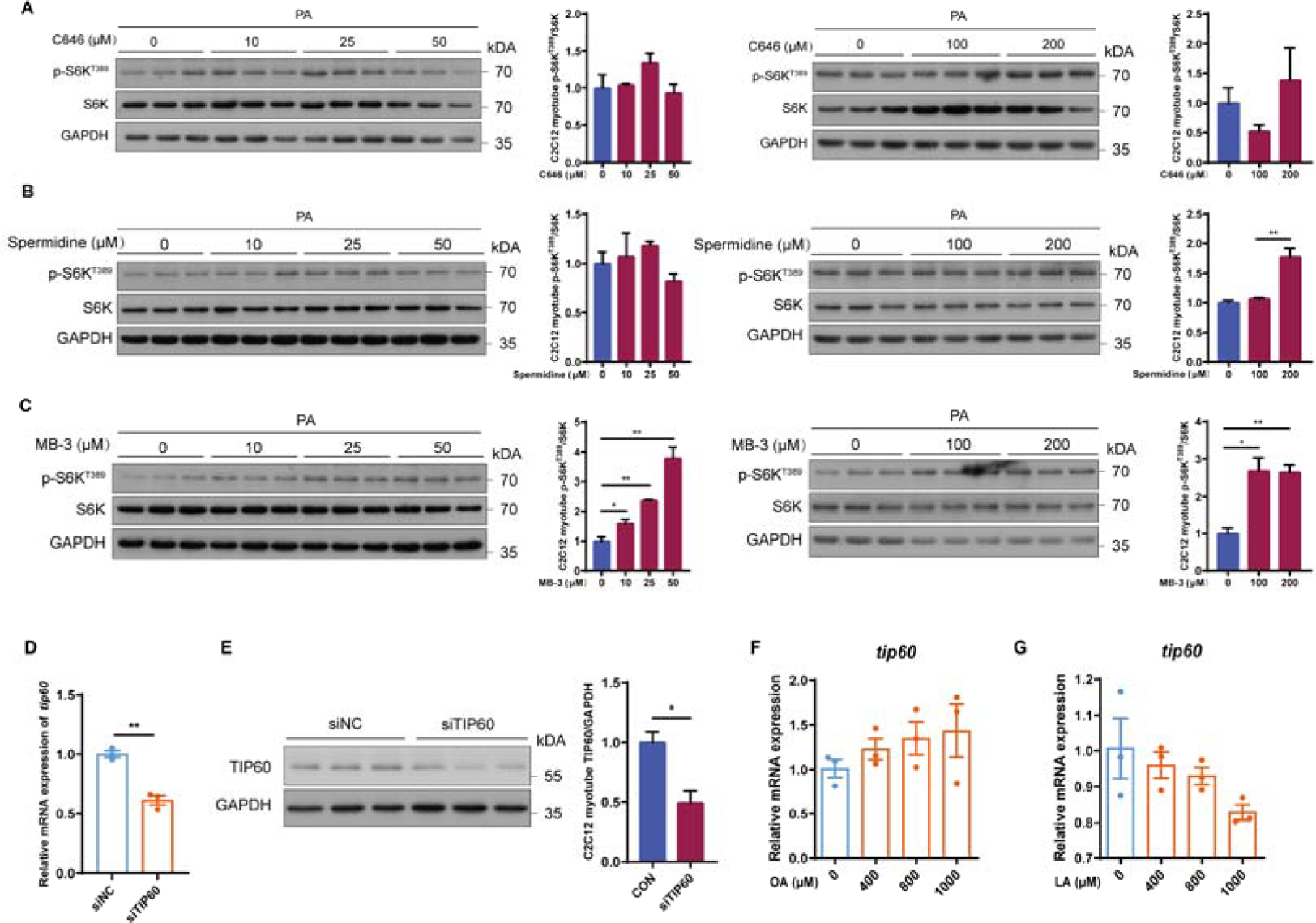
Tip60 regulates mTORC1 activity and insulin sensitivity through acetylating Rheb under PA treatment. Related to Figure 5. (A) The activity of mTORC1 signaling was measured by immunoblotting in C2C12 myotubes treated with the indicated concentrations of CBP/P300 inhibitor c646 under control or PA treatment for 12 h (n=3). (B) The activity of mTORC1 signaling was tested by immunoblotting in C2C12 myotubes treated with the indicated concentrations of CBP/P300 inhibitor spermidine under control or PA treatment for 12 h (n=3). (C) The activity of mTORC1 signaling was tested by immunoblotting in C2C12 myotubes treated with the indicated concentrations of GCN5 inhibitor MB-3 under control or PA treatment for 12 h (n=3). (D and E) Relative mRNA (D) and protein (E) levels of Tip60 were analyzed by quantitative PCR and immunoblotting in C2C12 myotubes transfected with control siRNA or siRNA against Tip60 for 48 h. (F) Relative mRNA levels of *tip60* were analyzed by quantitative PCR in fish myocytes under the indicated concentrations of OA treatment for 12 h (n=3). (G) Relative mRNA levels of *tip60* were detected by quantitative PCR in fish myocytes under the indicated concentrations of LA treatment for 12 h (n=3). The results are presented as the mean ± SEM and were analyzed using independent *t*-tests (\**p* < 0.05, \*\**p* < 0.01).

**Figure S6.**
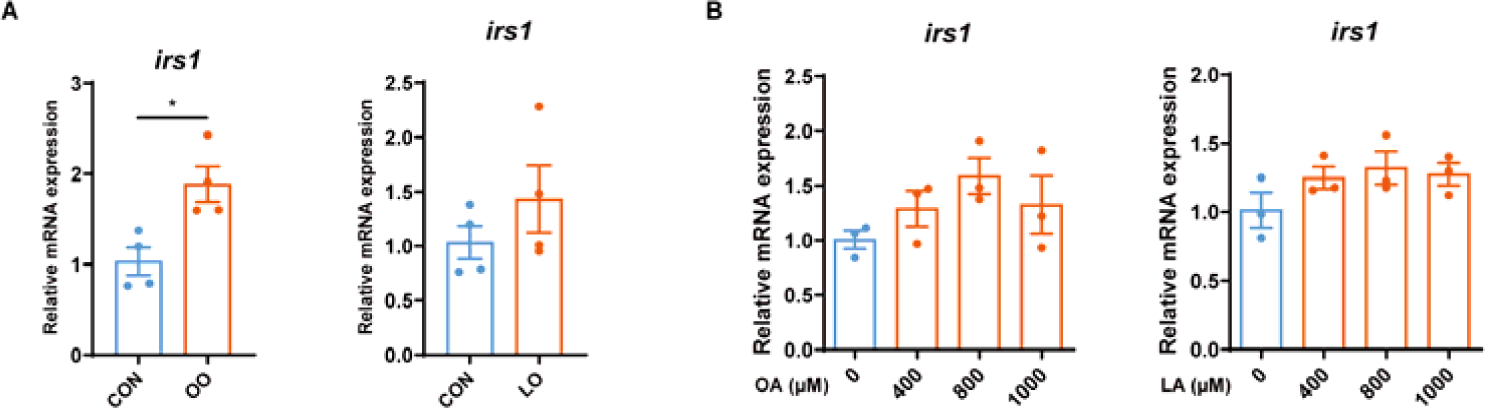
The effect of OA and LA on the relative mRNA levels of *irs1*. Related to Figure 6. (A) Relative mRNA levels of *irs1* were tested by quantitative PCR in the muscle of fish fed CON, OO or LO diet (n=4). (B) Relative mRNA levels of *irs1* were analyzed by quantitative PCR in C2C12 myotubes under the indicated concentrations of OA or LA treatment for 12 h (n=3). The results are presented as the mean ± SEM and were analyzed using independent *t*-tests (\**p* < 0.05).

**Table S1.** Fatty acid profiles of the experimental diets^a^.

| Fatty acid | CON | LO | OO | PO |
| --- | --- | --- | --- | --- |
| C14:0 | 8.08 | 2.06 | 1.97 | 1.64 |
| C16:0 (PA) | 36.88 | 27.21 | 28.76 | 56.95 |
| C18:0 | 6.69 | 7.03 | 5.26 | 5.07 |
| C20:0 | 0.16 | 0.26 | 0.23 | 0.07 |
| ΣSFA | 51.81 | 36.56 | 36.22 | 63.73 |
| C16:1n-7 | 3.63 | 0.74 | 0.85 | 0.48 |
| C18:1n-9 (OA) | 11.49 | 16.85 | 46.95 | 22.21 |
| ΣMUFA | 15.12 | 17.59 | 47.8 | 22.69 |
| C18:2n-6 (LA) | 9.97 | 32.58 | 10.58 | 11.27 |
| C20:4n-6 | 0.19 | 0.04 | 0.02 | 0.02 |
| Σn-6 PUFA | 10.16 | 32.62 | 10.6 | 11.29 |
| C18:3n-3 | 1.22 | 2.53 | 0.81 | 0.41 |
| C20:5n-3 (EPA) | 4.4 | 1.67 | 1.43 | 0.94 |
| C22:6n-3 (DHA) | 4.22 | 1.02 | 0.82 | 0.51 |
| Σn-3 PUFA | 9.84 | 5.22 | 3.06 | 1.86 |
| n-3/n-6PUFA | 0.97 | 0.16 | 0.29 | 0.16 |
| Σn-3LC-PUFA | 8.62 | 2.69 | 2.25 | 1.45 |

<sup>a</sup>Fatty acid content is expressed as % total fatty acids.
CON, control diet; LO, LA-rich diet; OO, OA-rich diet; PO, PA-rich diet; SFA, saturated fatty
acids; MUFA, mono-unsaturated fatty acids; n-6 PUFA, n-6 poly-unsaturated fatty acids; n-3
PUFA, n-3 poly-unsaturated fatty acids.

**Table S2.**
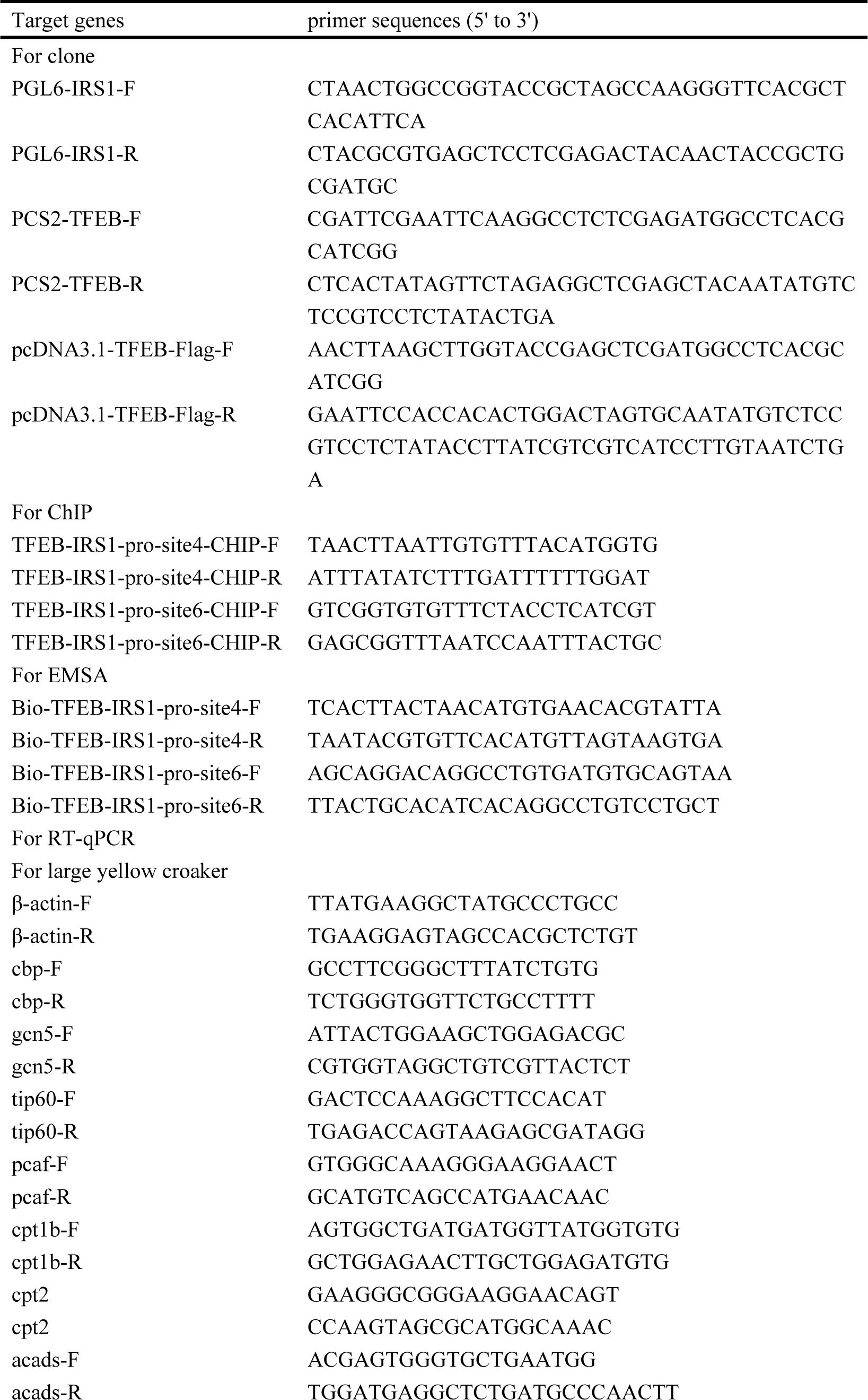
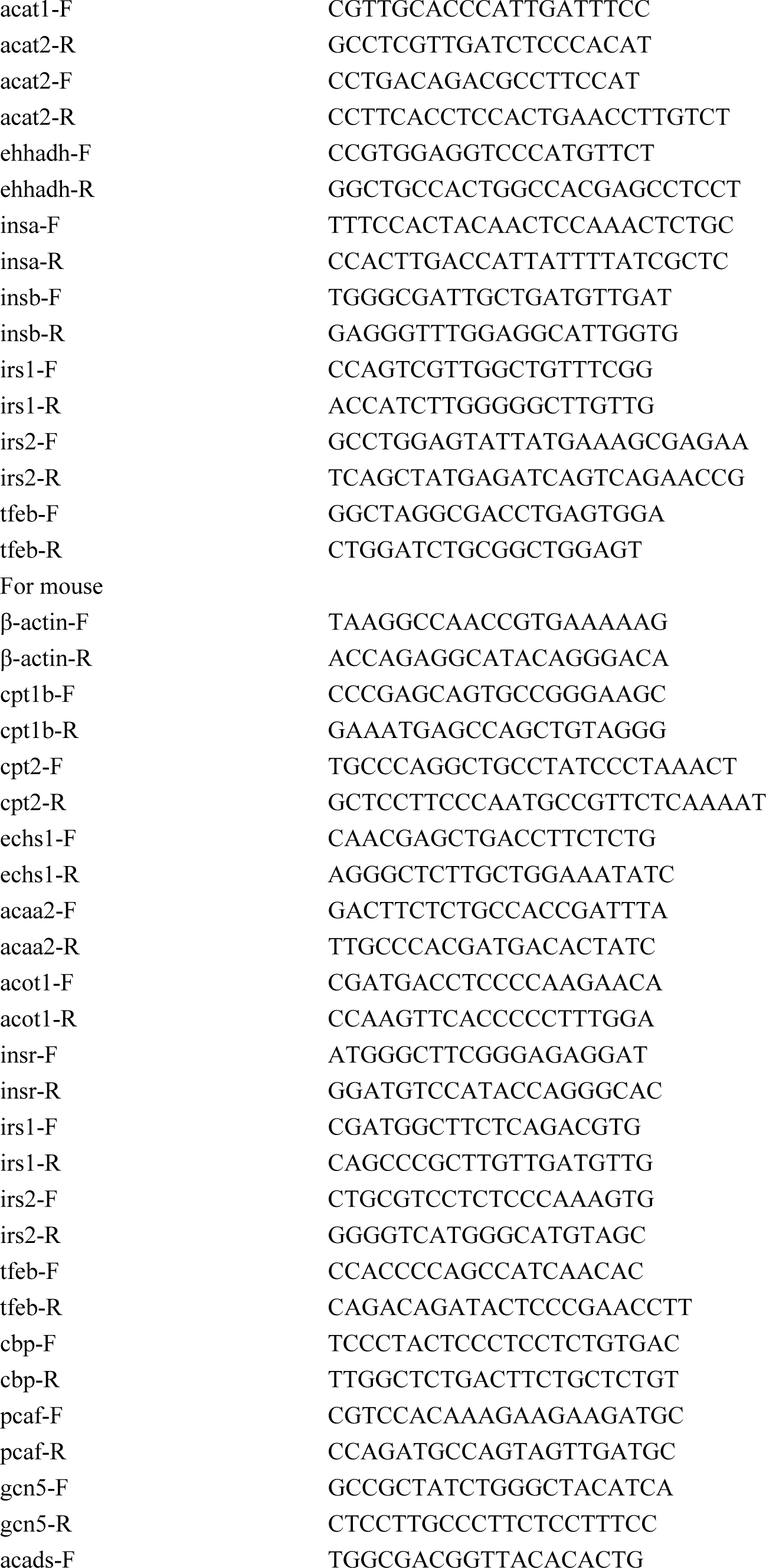
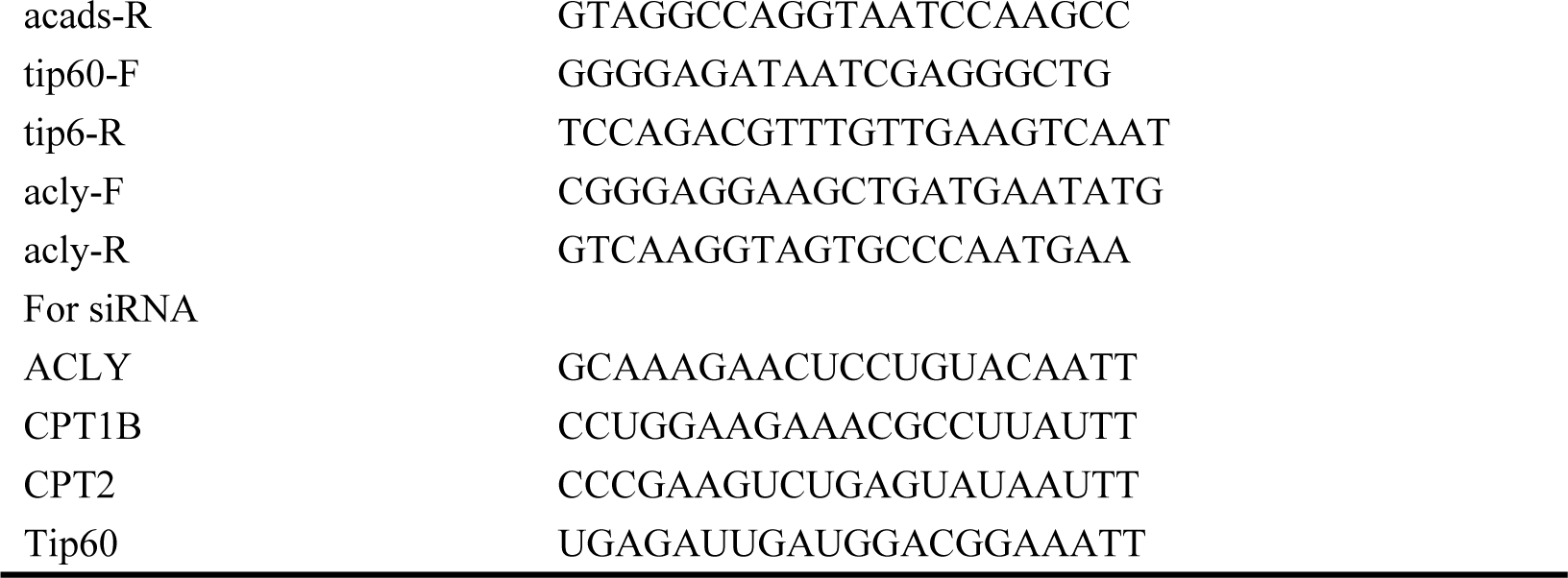
Sequences of the primers used in this study.

